# Investigating the differential structural organization and gene expression regulatory networks of lamin A Ig fold domain mutants of muscular dystrophy

**DOI:** 10.1101/2022.05.30.493956

**Authors:** Subarna Dutta, Vikas Kumar, Arnab Barua, Madavan Vasudevan

## Abstract

In metazoan cells, lamins form a proteinaceous meshwork as a major structural component of the nucleus and also play a role in regulating essential cellular processes. Lamins, along with their interactors, act as determinants for chromatin organization throughout the nucleus. The major dominant missense mutations responsible for autosomal dominant forms of muscular dystrophies reside in the Ig fold domain of lamin A. However, how lamin A contributes to the distribution of heterochromatin and balances euchromatin, and how it relocates epigenetic marks to shape chromatin states, remains poorly defined, making it difficult to draw conclusions about the prognosis of lamin A-mediated muscular dystrophies.In the first part of this report, we identified the in-vitro organization of full-length lamin A proteins due to two well-documented Ig LMNA mutations, R453W and W514R, through biophysical and electron microscopy observations. We further demonstrated that both lamin A/C mutant cells predominantly expressed nucleoplasmic aggregates with reduced amounts at the nuclear envelope. Labeling specific markers of epigenetics, such as H3K9me3, H3K27me3, H3K36me3, and HP1α, allowed correlation of lamin A mutations with epigenetic mechanisms. Immunofluorescence and biochemical analyses indicated transcriptional upregulation. In addition to manipulating epigenetic mechanisms, our proteomic studies traced diverse expressions of transcription regulators, RNA synthesis and processing proteins, protein translation components, and posttranslational modifications. These data suggest severe perturbations in targeting other proteins to the nucleus.Our study also predicts specific structural configurations of lamin A determined by its interacting partners’ patterns.

**Significance:**

- This study investigates the effects of two severe mutations in the LMNA gene associated with muscular dystrophies: **R453W** and **W514R**.
- This research aims to provide new insights into the structural environment of full-length lamin A protein due to single-point mutations.
- Our study employs proteomics to map expression levels of regulators involved in epigenetic states, RNA synthesis and processing, and protein synthesis and processing.
- It assesses potential effects on binding affinity and transcription, as well as alterations in heterochromatin protein distribution and RNA polymerase II functionality due to mutations affecting lamin A.
- A systematic investigation of lamin A interactors is conducted to explore their associations with other proteins.

**Graphical Abstract:** **Figure:** Schematic diagram representing the results. Mutations in lamin A protein evicts the nucleosome from nuclear periphery and consequently promotes aberrant gene expression.

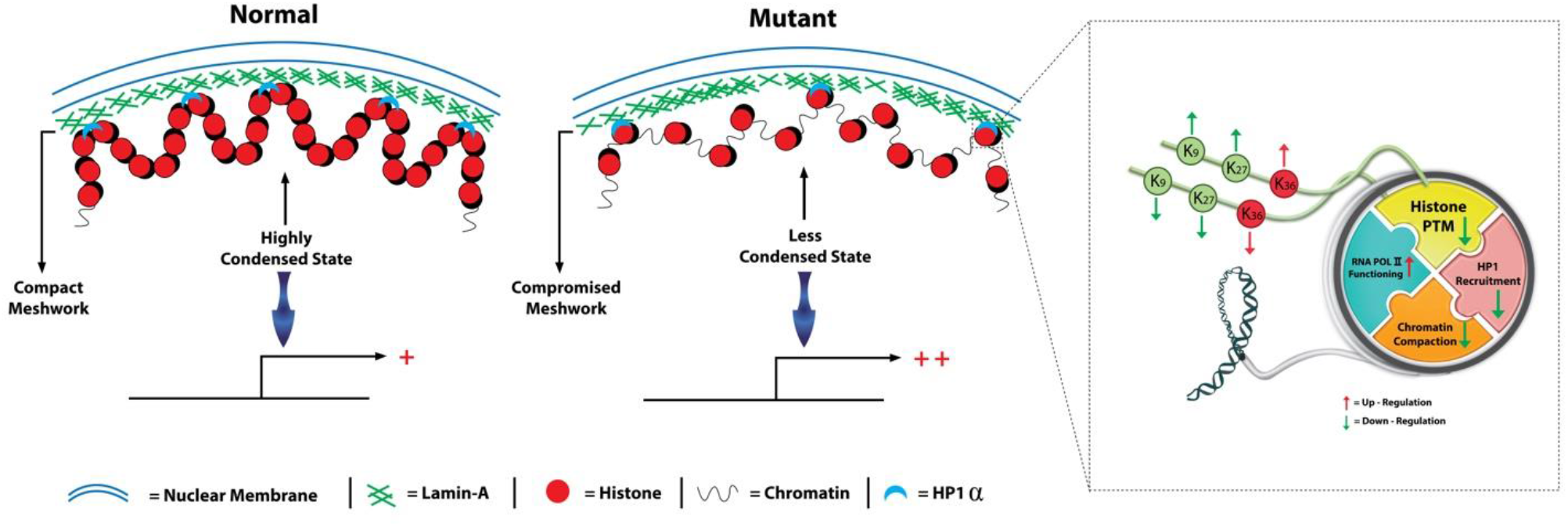

## Introduction

The nucleus is the most rigid entity of the cell, separated from the cytoplasm by a double membrane system known as the nuclear envelope (Dingwell C, 1992). This envelope consists of an outer nuclear membrane and an inner nuclear membrane, along with an array of nuclear pore complexes and a dense network of intermediate filaments called the lamina. The dense tetragonal networks of intermediate filaments are formed due to the separated but overlapping deposition of lamin proteins. In vertebrates, lamin genes express one A-type lamin and two B-type lamins, each with a molecular weight of 65-70 kDa. The prelamin forms of each type undergo post-translational modifications to form their mature forms (Adam et al., 2013; Maske et al., 2003; Dominici et al., 2009).

Lamins have a tripartite structure similar to type-V intermediate filaments, consisting of the N-terminal head domain (1-32 amino acids), the alpha-helical rod domain (33-383 amino acids), and the C-terminal globular tail domain (383-645 amino acids) (Bertrand AT et al., 2020; Strelkov SV et al., 2004; Dhe-Paganon S et al., 2002; Dutta S et al., 2018; Krimm I et al., 2002). The alpha-helical rod domain, the longest domain of the protein, exhibits a coiled-coil heptad-repeat pattern of three alpha-helical segments (coil 1A, coil 1B, and coil 2), interspersed with short intermediate sub-domains (L1 and L12) (Ahn et al., 2019). The C-terminal tail domain contains a distinct immunoglobulin-like (Ig) fold domain for chromatin binding, a nuclear localization signal (NLS) for lamin transport into the cell nucleus, and (except for lamin C) a cys-aliphatic-aliphatic-any residue box (CAAX) (Corrigan et al., 2005; Sinesky et al., 1994). The Ig-fold domain consists mainly of a compact β-sandwich with nine β-strands and short loops, arranged into two β-sheets (Dhe-Paganon et al., 2002). The rod domain and part of the short head region facilitate lamin assembly (Heitlinger et al., 1992; Isobe et al., 2007).

Parallel/unstaggered alpha-helical coiled-coil dimers of lamin A/C convert into high-order filamentous structures of 3.5 nm (distinct from intermediate filaments) with a repeating Ig-fold every 20 nm interval through lateral and longitudinal mixing (Koster et al., 2015; Parry et al., 2007; Turgay et al., 2017). Lamin A is particularly significant due to its involvement in over 500 autosomal dominant mutations associated with 16 tissue-specific degenerative diseases. Most of these mutations affect myopathies, with the Ig-fold domain bearing the highest number of mutations among laminopathies. Among mutations causing muscular dystrophy, 45% involve this domain. LMNA-associated muscular dystrophy is characterized by progressive skeletal muscle loss, skeletal and cardiac muscle weakness, cardiac dysrhythmias, and cardiomyopathy (Tan D et al., 2015).

Based on phenotypes such as herniated or misshapen nuclei and destabilized lamin structure, several hypotheses explain the pathogenesis of lamin A/C-associated diseases, focusing on cellular processes like nucleoskeleton-cytoskeleton coupling, protein-protein interactions, chromatin tethering, and transcription machinery. Two accepted hypotheses are the structural hypothesis, which emphasizes lamin’s mechanical integrity in maintaining nuclear homeostasis, and the gene regulation hypothesis, which highlights chromatin compartmentalization and epigenetic regulation of gene expression in tissue-specific lamin A/C diseases.

Lamin A plays significant roles in nuclear processes such as chromatin organization and positioning, DNA replication, transcription, and gene expression (Dechat et al., 2008; Malhas et al., 2007; Tang et al., 2008). Lamin B1 interacts closely with chromatin along the nuclear periphery, whereas lamin A cross-links chromatin throughout the nucleus, directly or indirectly through histone proteins (Melcer S et al., 2012; Prokocimer M et al., 2009; Moir RD et al., 2000). LMNA knockout results in chromatin rearrangement and deformation of the nuclear lamina. A-type lamins are responsible for gene relocation to the nuclear periphery, which may influence transcriptional pathways. Studies using DNA adenine methyltransferase identification (DamID) have identified genomic regions associated with the nuclear lamina, termed Lamina Associated Domains (LADs) (Guelen et al., 2008; Meuleman et al., 2013; Peric-Hupkes et al., 2010; Pickersgill et al., 2006; van Steensel and Kind, 2014; Kind et al., 2015). These regions are characterized by repressive histone post-translational modifications (PTMs), gene-poor content, and low transcriptional activity (de Laat W et al., 2008; Yao J et al., 2017; van Steensel B et al., 2013).

Lamin-binding proteins and DNA-bridging proteins, such as LBR, Emerin, BAF, histones, LAP2α, LAP2β, facilitate the connection between chromatin and the nuclear lamina (Mattout-Drubezki and Gruenbaum, 2003; Dorner et al., 2007; Schirmer and Foisner, 2007; Worman et al., 1990; Ye and Worman, 1994, 1996; Foisner and Gerace, 1993; Furukawa et al., 1997; Schumaker et al., 2001; Holaska et al., 2003). Lamin A/C regulates histone methyltransferases by modulating pRb signaling pathways (Djabali et al., 2010; Bracken et al., 2003; Nielsen et al., 2001; Dorner et al., 2007). Analysis of LMNA mutations in patients with familial partial lipodystrophy (FPLD), Hutchinson-Gilford progeria syndrome (HGPS), and autosomal dominant Emery-Dreifuss muscular dystrophy (AD-EDMD) reveals significant anomalies in peripheral heterochromatin (Sabatelli et al., 2001; Capanni et al., 2003; Filesi et al., 2005; Goldman et al., 2004; Columbaro et al., 2005). Recent reports highlight the influence of distinct epigenetic landscape patterns on heterochromatic changes in mutated and lamin A null cells (Gonzalez Suarez et al., 2009; Shumaker et al.2006).

Constitutive heterochromatin is characterized by trimethylation of histone H3 at Lys9, histone H4 at Lys20 (H4K20me3), and histone H3 at Lys27 (H3K27me3), while facultative heterochromatin is marked by histone H3 at Lys27 (H3K27me3) (Sarma and Reinberg, 2005; Martin and Zhang, 2005). Lamin A/C deficiency leads to a loss of the heterochromatin-euchromatin ratio, along with rearrangement of HP1β and H3K9 methylation (Filesi et al., 2005; Galiova et al., 2008). In HGPS fibroblasts, downregulation of HP1α, dislocalization from H3K9me3 sites, and decreased levels of H3K9me3 and H3K27me3 are observed, alongside upregulation of H4K20me3 (Scaffidi and Misteli, 2005; Shumaker et al., 2006; Columbaro et al., 2005). Older individuals expressing LAΔ50/progerin and mandibuloacral dysplasia fibroblasts exhibit anomalies in H3K9me3 levels and loss of H3K27me3 (Filesi et al., 2005; Scaffidi and Misteli, 2006). Overexpression of wild-type lamin A reduces H3K4 methylation in C2C12 cells, and muscular dystrophy cells (R453W lamin mutation) lose H3K4 hypermethylation at the MYOG promoter (Collas P et al., 2008).

Numerous studies support the role of lamin A in transcriptional regulation, where condensed chromatin structure affects transcription initiation by RNA Pol II (Misteli T et al., 2009; Workman JL et al., 2007). A dominant negative construct of lamin A perturbs RNA polymerase II and the localization of the TATA binding protein (TBP) (Kumaran et al., 2002). Transfection of the ΔNLA mutant (deletion of the N-terminal domain) into BHK 21 cells reduces BrUTP incorporation, indicating dysregulation in transcription (Spann et al., 2002). Conversely, overexpression of lamin A/C suppresses RNA Pol II activity in HeLa cells (Kumaran et al., 2002). These findings underscore the impact of lamin A on chromatin organization and regulation, spanning from gene expression to transcriptional activity. Furthermore, transcriptional activity correlates with RNA metabolism and protein metabolism (McCracken S et al., 1997; Shuman S et al., 1997; Hirose Y et al., 1998). Interestingly, HGPS-derived fibroblasts exhibit enhanced RNA processing, including mRNA splicing via the spliceosome, RNA transport, protein translation, and ribosome biogenesis (Buchwalter A et al., 2017). Splicing factors have been found associated with lamin A (Spann et al., 2002; Kumaran et al., 2002; Depreux FF et al., 2015).

Our study investigates the epigenetic mechanisms underlying chromatin organization and the proteome profile of the global gene expression regulatory network in lamin A-associated muscular dystrophy. We focused on two severe mutations of muscular dystrophy, that reside in Ig fold domain of LMNA : R453W and W514R. According to the database(www.umd.be/LMNA), 66 records have reported the occurrence of the R453W mutation in exon 7 of the LMNA gene, associated with Emery-Dreifuss Muscular Dystrophy (EDMD). On the other hand, 3 records have cited the W514R mutation in exon 9 of the LMNA gene, implicated in skeletal muscular dystrophy. Krimm et al.2002 have previously reported significant destabilization in the Ig structure caused by the R453W mutation, potentially impacting its differentiation capability, as noted by Favreau et al.2004.However, no comprehensive structural analysis of full length R453W LA,W514R LA mutation have been reported to date. Dialynas et al.2015 conducted a basic examination of tertiary structure changes using 15N/1H HSQC NMR for the W514R mutation. The complete molecular mechanism underlying the pathogenesis of these muscular dystrophies remains unknown.

Using a proteomics approach, we mapped the expression levels of regulators involved in epigenetic states, RNA synthesis and processing, and protein synthesis and processing, respectively. Additionally, we conducted a systematic investigation of lamin A interactors to gain new insights into their associations with proteins.This study is designed to provide additional insights into the structural environment of the full-length lamin A protein due to single-point mutations in the Ig fold domain, and their potential effects on binding affinity and transcription. Our results also support alterations in heterochromatin protein distribution and the functioning of RNA polymerase II, offering predictions about gene expression patterns.

## Results

### Mutation affects gel like behaviour of lamin A and the secondary structure

Lamin provides substantial support to the nucleus, as lamin A deficiency results in irreversible fluid-like deformation (Pajerowski et al., 2007). Mutations in the LMNA gene disrupt nuclear compliance and reduce chromatin stability, with effects varying based on specific amino acid substitutions (Zwerger et al., 2013; Earle et al., 2020). Recent studies by Banerjee et al. (2013) and Mukherjee et al. (2021) have highlighted differential rheological properties associated with different point mutations linked to dilated cardiomyopathy. Therefore, it is plausible to speculate that Ig-fold mutations used in this study exhibit variable elastic properties. Proteins were purified to >95% homogeneity using a cation exchange column (Figure S1). For biophysical experiments, proteins of similar purity were used. During renaturation with stepwise urea removal, R453W exhibited maximal precipitation. Equal amounts of proteins were blotted with anti-lamin A antibody, showing robust, clean bands. The high homogeneity of the preparation ensures that the results reflect the protein of interest and not impurities.

We tested the effects of constant oscillatory shear strain amplitude (γ) of 1% at an angular frequency (ω) of 5 rad/s for 1000 seconds on the viscoelastic properties of wild-type and mutant proteins at a concentration of 0.6 mg/ml (Figure 1-b). W514R lamin demonstrated a more solid-like nature compared to wild-type lamin, evidenced by higher storage moduli (G’). In contrast, R453W lamin consistently showed lower storage moduli (G’) than the native protein. Interestingly, we observed higher saturation levels in both storage moduli (G’) and loss moduli (G”) for wild-type lamin, confirming stable tertiary structure. The different values of storage moduli (G’) for R453W lamin and W514R lamin indicate disparities in polymerization population, load support, and force transmission mechanisms.

**Figure 1.**
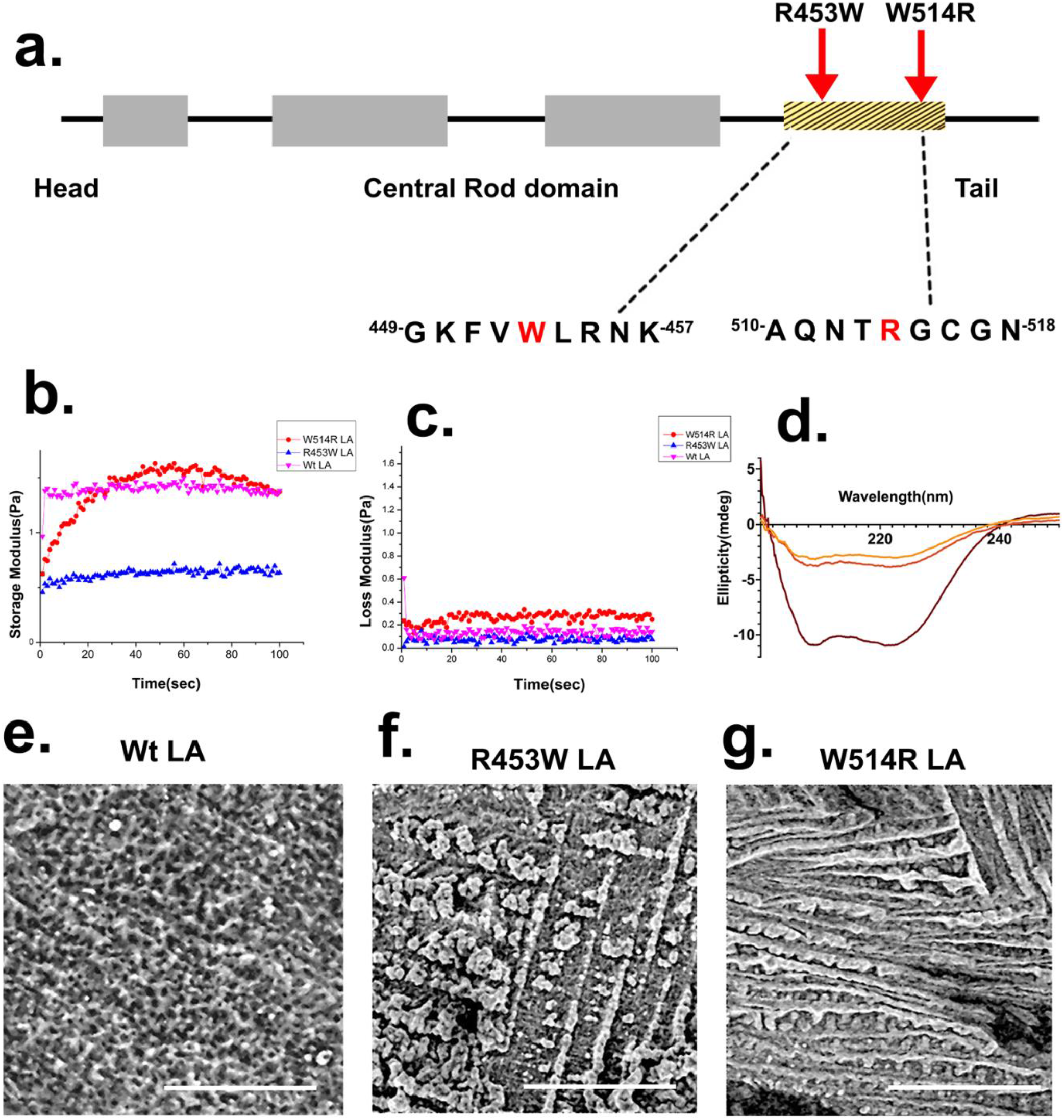
Structural perturbation due to lamin A mutation. (a) Represents position of twomutations in Ig fold of lamin A. (b),(c) Viscoelastic spectra (G’> G”) of lamin A changed due to mutations. (d) Far-UV CD spectra of 4 μM LA shows secondary structure of LA is altered by point mutations. (e),(f),(g) Differential bundle –like networks and crisscross density in musculardystrophy causing mutants compared to Wt LA by scanning electron microscopy. Scale bar = 20μm, magnification 3000x.

The variability in viscoelastic properties likely reflects differences in lamin A assembly patterns. Our scanning electron microscopy data (Figure 1-c) showed that W514R lamin tends to form bundled filaments with fewer visible cross-links compared to the densely cross-linked network observed with wild-type lamin, which directly influences protein strain hardening behavior. Conversely, R453W lamin exhibited weak network formation with fewer bundles and cross-links. Anomalies in the criss-cross density of mutant proteins correlated well with complex viscosity plots, which indicated higher viscosity for W514R lamin compared to wild-type lamin A, and lower viscosity for R453W lamin.

Changes in viscoelastic properties and assembly patterns due to mutations can be understood by examining the secondary structure of proteins, particularly the behavior of the rod domain. Interestingly, our circular dichroism spectroscopy analysis revealed non-overlapping spectra among the mutants and the native protein. These alterations also suggest differences in the oligomerization status of the proteins, with mutants inducing elevated magnitudes of [*θ*] in their oligomerized forms compared to the wild-type, indicating enhanced and/or altered homopolymerization. Specifically, we calculated that the *θ*222/*θ*208 ratio increased to approximately 1.2 for W514R, confirming increased coiled-coil formation due to self-association of mutant lamin A monomers; whereas for R453W, this ratio decreased to approximately 0.89, suggesting varying degrees of alteration in lamin A helicity due to mutations.

The *θ*222/*θ*208 ratios are displayed in **Table 1**. A ratio exceeding 1 defines the presence of coiled-coils.

**Table 1:**

| **Proteins** | **$[\theta]_{222}/[\theta]_{208}$** |
| --- | --- |
| **Wt LA** | **.91** |
| **R453W LA** | **.89** |
| **W514R LA** | **1.2** |

The impact of mutation on the thermodynamic stability of the protein was initially assessed using DynaMut2. The server predicted the change in the free energy of unfolding (ΔΔG) induced by the mutations. For instance, the mutation Arg453Trp resulted in a destabilizing effect with a negative ΔΔG of -0.8 kcal/mol (Figure S2). Conversely, Trp514Arg was found to be mildly stabilizing, showing a positive ΔΔG of 0.23 kcal/mol, suggesting the removal of a configurational constraint. Interestingly, different ΔΔG values were obtained for mutations affecting chain A individually, indicating that the environments of these mutations within the A chain are dissimilar.

### Nuclear architecture analysis by 3D SIM and mathematical modelling

Previous investigations into lamin A organization within mammalian cell nuclei have revealed structural disparities between wild-type and mutant lamin A cells (Dutta et al., 2018). In this study, we extended our comprehension of nuclear landscapes using three-dimensional structured illumination microscopy. We identified significant insights into the nanoscale distribution of lamin A within stably transfected wild-type lamin A cells(Figure 2-a). Our findings illustrate the formation of a network-like structure throughout the nucleus, accompanied by prominent rim staining. Figure 2-b,c further illustrates how these architectural features are altered in mutant lamin A cells. Notably, increased mesh size, disrupted rim staining, and the presence of aggregates were observed in nuclei expressing R453W and W514R lamin A. The enlargements in laminar mesh size are depicted in Figures 2-d, 2-e, and 2-f. These results indicate that abnormal self-assembly of lamin A is responsible for aggregate formation, stemming from inherent structural modifications in LMNA proteins.

**Figure 2.**
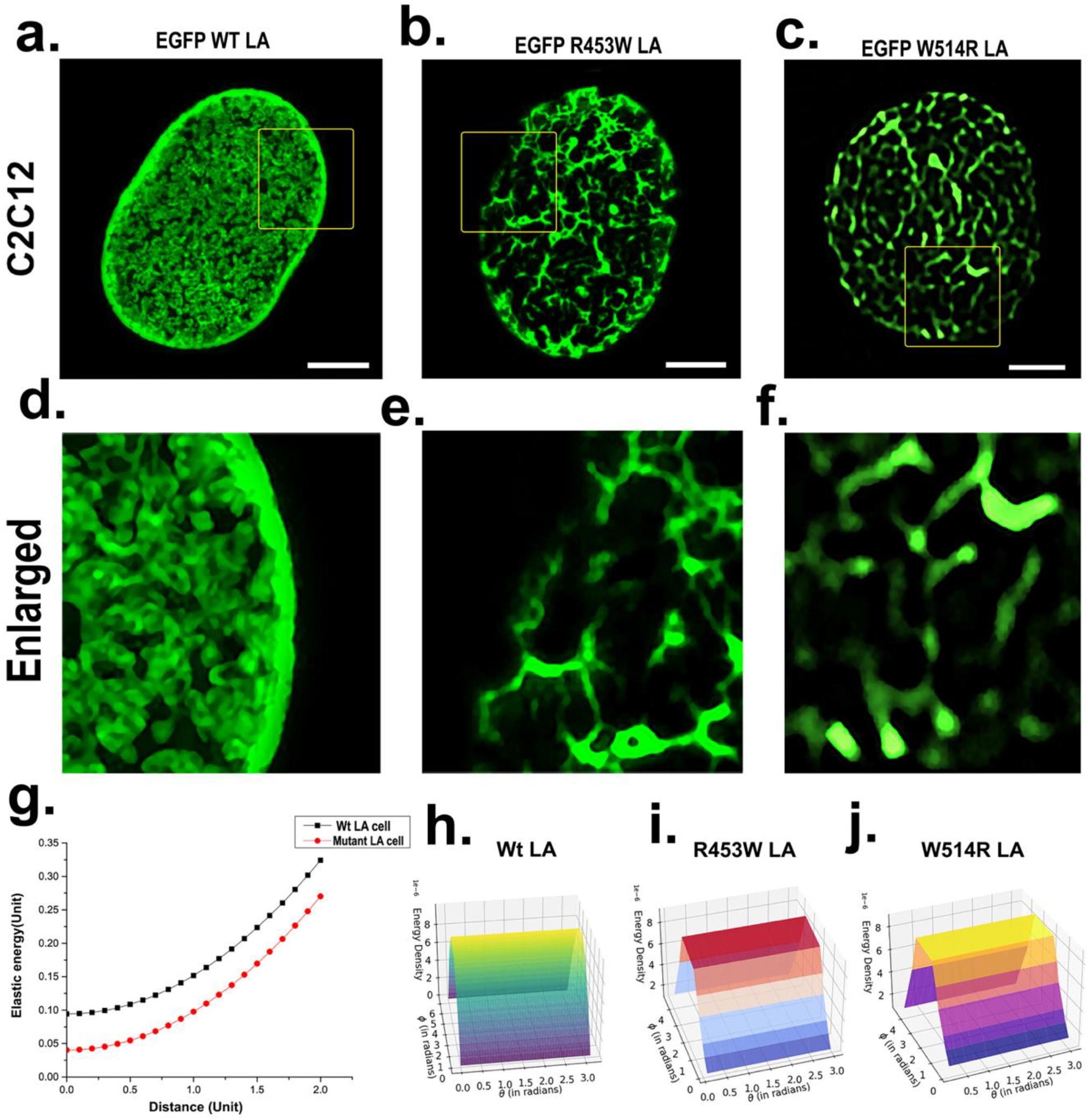
Aberrant laminar organization in cells expressing muscular dystrophy mutants of LA. (a),(b),(c) N-SIM micrographs of LA meshwork in Wt and R453W, W514R expressing C2C12 cells; bars 2 μm. (d),(e),(f) panels represent enlarged nuclear meshwork marked by yellow squares in the upper panel. (g),(h),(i) and (j) Representative elastic energy profiles for Wt/mutant LA. For Wt LA cell (*θ*=0 (300 nodes), *θ*=*π*/10 (250 nodes), *θ*=2*π*/10 (200 nodes), *θ*=3*π*/10 (150 nodes), *θ*=4*π*/10 (100 nodes), *θ*=5*π*/10=*pi*/2=(50 nodes)(black line)), 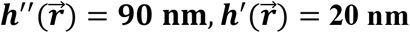 and for Mutant LA cell (*θ*=0 (50 nodes), *θ*=*π*/10 (45 nodes), *θ*=2*π*/10 (40 nodes), *θ*=3*π*/10 (35 nodes), *θ*=4*π*/10 (30 nodes), *θ*=5*π*/10=*π*/2=(25 nodes) (red line)),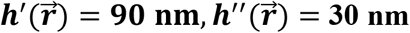.

To delve deeper into the impact of changes in the mesh size of the nucleus, which can potentially affect its deformability, we aim to compute elastic energy through mathematical modelling. Elastic energy refers to the mechanical potential energy stored within a material or system due to the deformation of its volume or shape. In our analysis, this involves calculating the strain tensor, which quantifies the deformation or displacement within a continuous medium and is directly proportional to strain. Our model assumes the nucleus comprises nuclear envelope proteins and nuclear lamin A, and we treat it as a thin spherical shell for these calculations. Our mathematical modeling-based framework demonstrates changes in elastic energy that are influenced by lamin A mutations. Figure 2-g,h,i and j indicate variations in elastic energy.

### Alteration of histone modification marks as a sequel to lamin A mutations

Several experimental studies have suggested that histone modification marks play a role in lamin A mutations (Lattanzi et al., 2007; Scaffidi and Misteli, 2006; Shumaker et al., 2006; Filesi et al., 2005). In our immunofluorescence study, we examined H3K9me3 and H3K27me3 as markers of transcriptional activation, and H3K36me3 as a marker of repression. H3K9me3 and H3K27me3 are typically associated with heterochromatin (Kouzarides et al., 2007), while H3K36me3 is associated with euchromatin (Bartova et al., 2008; Berger, 2007).

We observed a significant decrease in H3K9me3 dots, with more pronounced reduction in EGFP LA-R453W compared to EGFP LA-W514R (Figure 3). Unexpectedly, histone marks were found to colocalize with aggregates in the R453W mutation, consistent with the known propensity of both R453W and W514R lamin A mutants to form aggregates (Roblek, Ogris et al., 2010; Dutta et al., 2018). Specifically, larger globular dots of H3K27me3 marks were observed in R453W lamin A, albeit at lower counts, while a diffuse staining pattern of H3K27me3 marks was noted in LA W514R. The presence of high accumulation of H3K36me3 dots in the nucleus of mutants further supported these findings. Figure 3 illustrates these observations.

**Figure 3.**
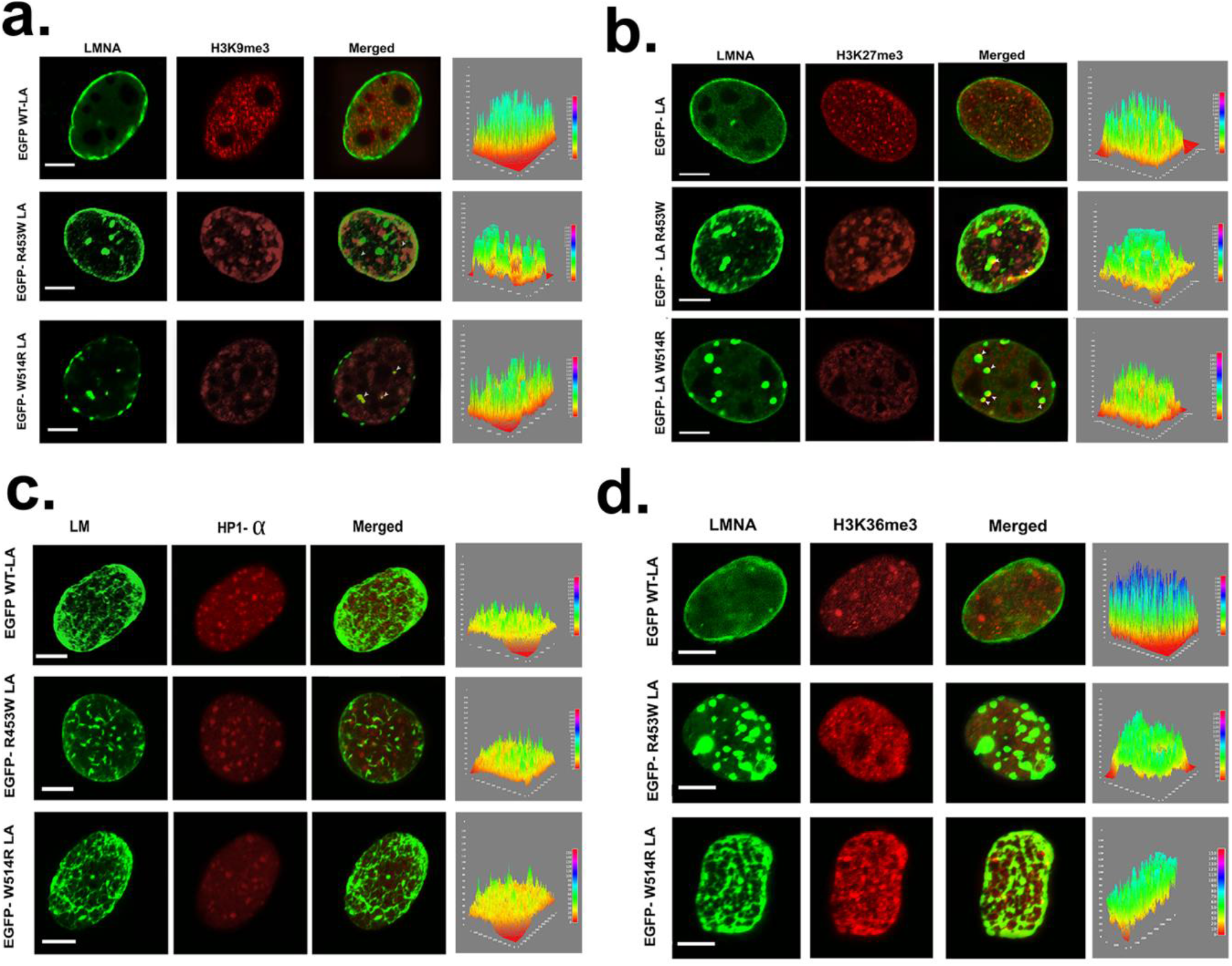
Alteration of repressive epigenetic marks : H3K9 me3 (a), H3K27me3 (b), HP1α (c) and active epigenetic mark : H3K36me3 (d). Stably transfected C2C12 cells with pEGFP WT LA, pEGFP R453W LA, pEGFP W514R LA were immunostained with anti-H3K9me3,anti-H3K27me3, anti-HP1α, anti-H3K36me3 following counterstaining with Alexa fluor 568. 3D surface plots of each red channel from the image J represent the distribution of spatial fluorescence intensity of each samples. The X axis features the distance along the line and the Y axis is pixel intensity.

Immunoblot analysis of whole cell lysates expressing EGFP WT LA and mutants (Figure 5) confirmed the results obtained from immunofluorescence studies, showing decreased expression levels of H3K9me3 and H3K27me3, and increased expression of H3K36me3 in the mutants. The original blots are presented in Figure S3.

**Figure 4.**
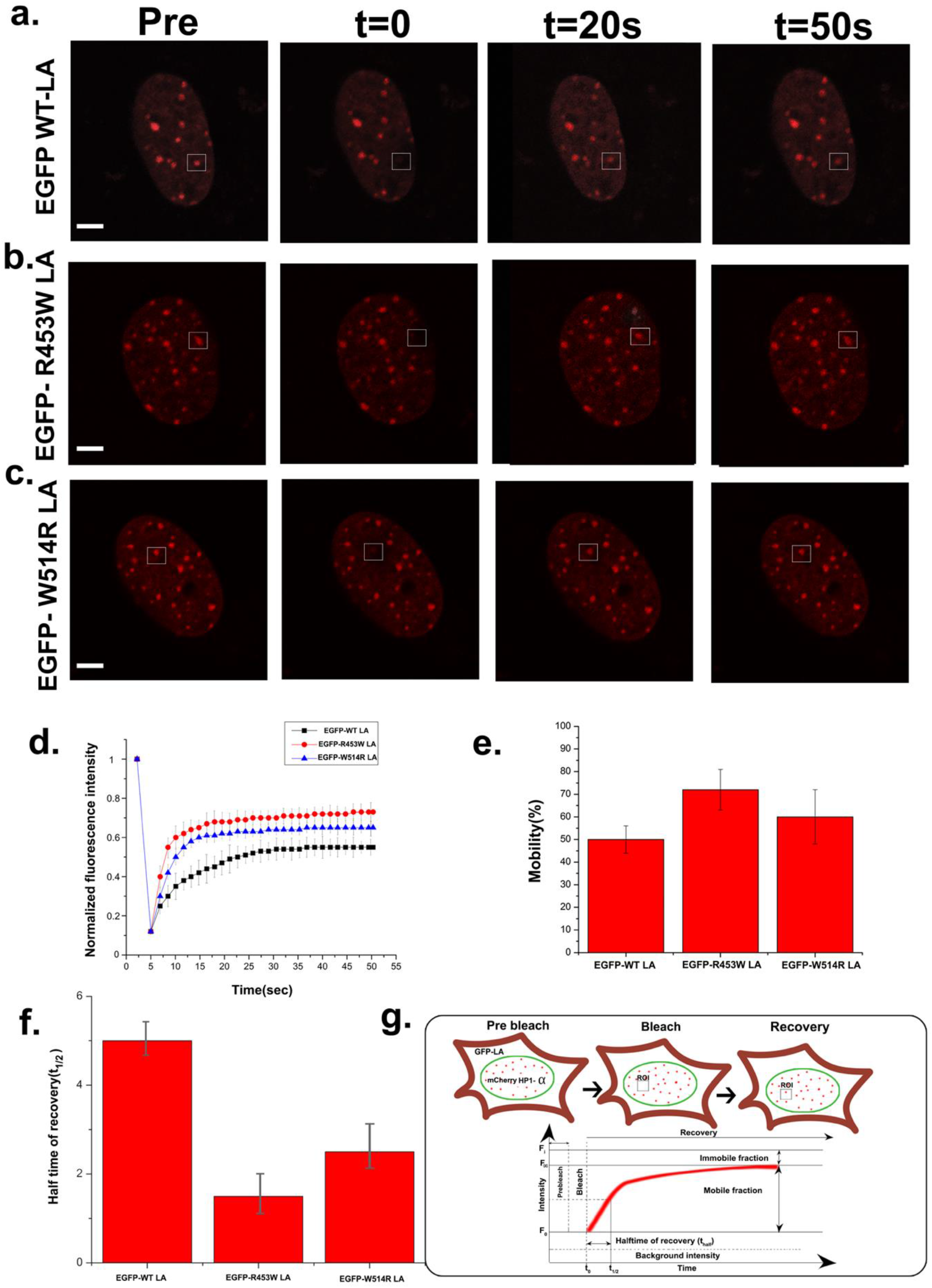
Alteration of HP1α dynamics in mutant lamin A. (a). Representative time series FRAP images of RFP-HP1α immediately following the photo bleach (t=0), Bar,10µm. C2C12 cells were transiently transfected with EGFP-WT / EGFP-R453W / EGFP W514R –RFP HP1α for 24 hours prior to FRAP. (b). FRAP analysis of RFP HP1α in wild type and mutants. Y axis denotes normalized florescence intensity which points percentage of recovery. (c) Half time of recovery (t1/2) graph is generated for all the samples from half maximal fluorescence recovery (F1/2). (d). Mobility graph is generated from the data obtained from normalized FRAP curves for all the samples. Data show the mean ± standard deviation for 30 cells of each sample.

**Figure 5.**
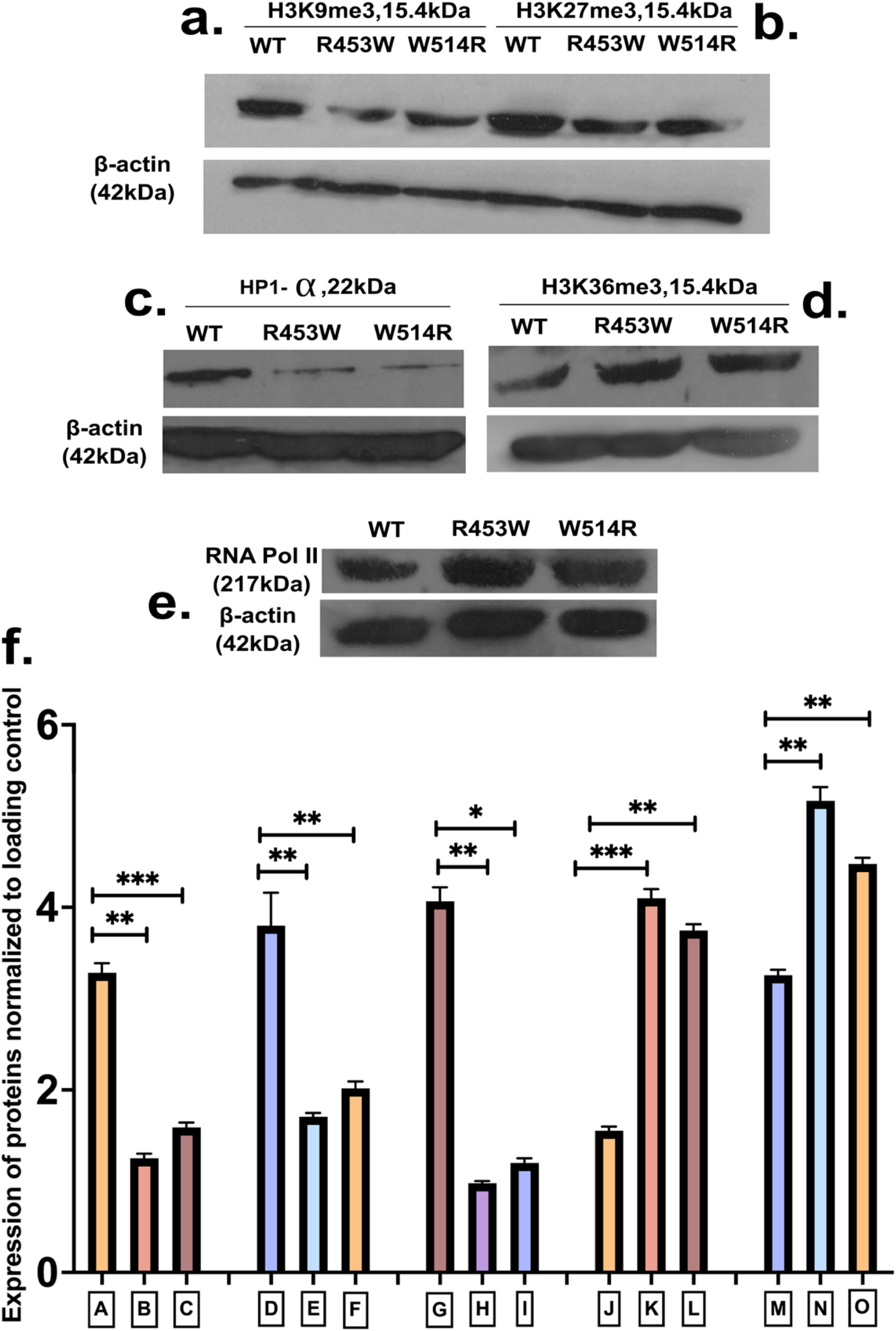
Immunoblot analyses of H3K9me3, H3K27me3, H3K36me3, HP1α, RNA Pol II expression in C2C12 cells. Cell lysates from stably transfected cells run in SDS PAGE and immunoblot with anti-H3K9me3(a), anti-H3K27me3(b), anti-H3K36me3(d), anti-HP1α(c), anti-RNA Pol II(e) respectively, β-actin was used as a loading control. Quantification of the expression levels shown in (f).

### Alteration of HP1 α and dynamics in mutant lamin A cell

Depletion of HP1 proteins has been shown to alter the genomic landscape by reducing H3K9 methylation marks (Maeda et al., 2022). High levels of HP1α are characteristic of constitutive heterochromatin (Trojer and Reinberg, 2007). In our immunofluorescence studies, we observed low recruitment of HP1α in heterochromatin foci for mutant lamin A, confirming the reduced levels of H3K9me3 observed in both our immunofluorescence and immunoblot analyses (Figure 3-c). These findings were consistent with the immunoblot results, which demonstrated decreased intensity levels for both LA R453W and LA W514R (Figure 5-c). Our observations align with previous studies that have reported HP1α depletion in lamin A rod mutants (Chaturvedi, Parnaik, 2010).

Furthermore, our FRAP (Fluorescence Recovery After Photobleaching) experiment (Figure 3) indicated increased dynamics of HP1α due to mutant lamin A. This suggests enhanced oligomeric exchange between protein molecules, resulting in less stable complex formation with H3K9me3 and reduced recruitment of HP1α to heterochromatin. Importantly, our calculations (also detailed in Figure 4-f) yielded half-time (t1/2) of recovery values of 5 ± 0.625 s for EGFP WT LA, 1.8 ± 0.225 s for EGFP R453W LA, and 2.5 ± 0.312 s for EGFP W514R LA.

### Hyperactivation of RNA pol II functionality

Several studies have suggested that lamins play roles in RNA polymerase II (RNA Pol II) activity, either directly or indirectly (Timothy Spann, Robert D. Goldman, 2002; Kumaran Ri et al., 2002; Hessen and Fornerod, 2007). The activity of RNA Pol II has been shown to be disrupted in cases involving dominant negative lamin A with terminal deletions (Spector DL et al., 2008). Phosphorylation of serine residues in the C-terminal domain (CTD) of RNA Pol II at positions 2, 5, and 7 is crucial for its activity, particularly Serine 5 phosphorylation which is associated with transcription initiation (Chapman RD et al., 2008).

According to immunoblot data, LA R453W showed significantly higher levels of RNA Pol II Serine 5 phosphorylation (RNA Pol II s5p) compared to LA W514R and wild type lamin A (Figure 5).This suggests that mutations in lamin A, such as R453W and W514R, may influence RNA Pol II activity differently, potentially impacting transcription initiation processes.

### Proteome profile of LA mutants

Total proteome profiling of LA R453W and LA W514R mutants was conducted in triplicates, with wild type samples analyzed in duplicates. ProteinPilot™ Software 5.0 (AB Sciex) was employed for protein searches against the UniProt mouse database. The search criteria included an unused value (ProtScore) threshold of 1.3, and proteins identified with a confidence level of 95% or higher were included in the analysis from the LC-MS/MS data. A total of 952 proteins were resolved in at least one of the eight samples.

Given the presence of replicates for both mutants and wild type, Principal Component Analysis (PCA) and Euclidean-based hierarchical clustering were utilized to assess the maximum variation across all samples and replicates (Figure 6). PCA was plotted based on the total number of proteins expressed and their expression levels to evaluate the consistency of biological replicates. Closer proximity on the PCA plot indicates similar expression profiles among samples. The hierarchical clustering tree was constructed using a distance metric that considers both the number of proteins expressed and their respective expression levels. Replicate samples clustered closely together, indicating minimal variation among them.

**Figure 6.**
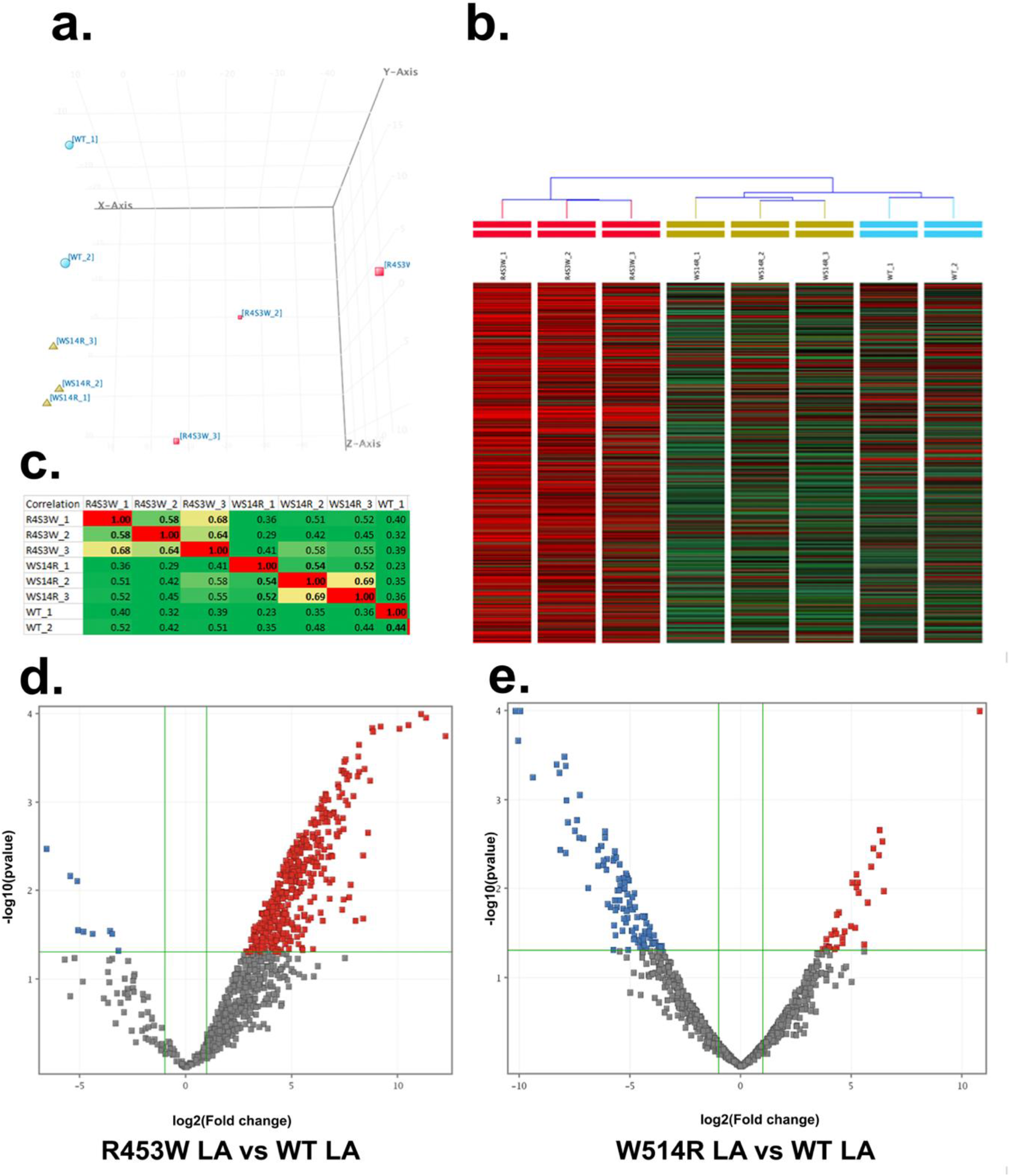
Qualitative and quantitative assessment of proteome profile: (a), (b), (c) PCA analysis, unsupervised condition tree, correlation plot presented reproducibility among 952 proteins in at least one of the eight samples. (d),(e) volcano plots of R453W LA, W514R LA respectively. Out of 952 proteins, upregulation of 860 proteins and downregulation of 91 proteins for LA R453W and upregulation of 392 proteins and downregulation of 559 proteins for LA W514R were observed.

Differential expression analysis was performed using an Anova t-Test with a significance threshold (p-value) of 0.05 and a fold change criterion of ≥2 for proteins to be considered differentially expressed. Visualizing the results with a volcano plot revealed that LA R453W showed upregulation of 860 proteins and downregulation of 91 proteins, whereas LA W514R exhibited upregulation of 392 proteins and downregulation of 559 proteins compared to the wild type. Unsupervised hierarchical clustering further illustrated distinct patterns of upregulated and downregulated proteins in both mutants relative to the wild type sample.

### Quantitative assessment on gene expression regulatory proteins

For the differentially expressed proteome analysis, comprehensive biological functional annotations of the identified proteins were conducted using the UniProt database website and DAVID data analysis software (http://david.abcc.ncifcrf.gov/home.jsp). The analysis encompassed Gene Ontology (GO) categories such as biological processes, cellular components, and molecular functions to provide an overall distribution of the differentially expressed proteins. These proteins were notably involved in various pathways including epigenetic regulation, RNA synthesis and processing, protein synthesis and processing, and ribosomal functions(Figure S4).

To further explore the regulatory networks influenced by the mutations, we performed pathway enrichment analysis using in-house scripts and visualized the results using Cytoscape V2.8.3. This analysis helped identify key nodes and edges of gene expression regulatory proteins that were uniquely dysregulated by both mutants compared to the wild type (Figure 7). Significant functional categories and pathways were determined using hypergeometric test-derived p-values, corrected for false discovery rate (FDR) with a threshold of 0.05.In our study, we observed upregulation of 272 proteins and downregulation of 25 proteins in the R453W LA sample, while the W514R LA sample showed 141 upregulated proteins and 57 downregulated proteins across all selected subgroups. Notably, regulatory proteins involved in protein synthesis and processing were found to be downregulated (36 proteins) specifically in the W514R LA sample.

**Figure 7.**
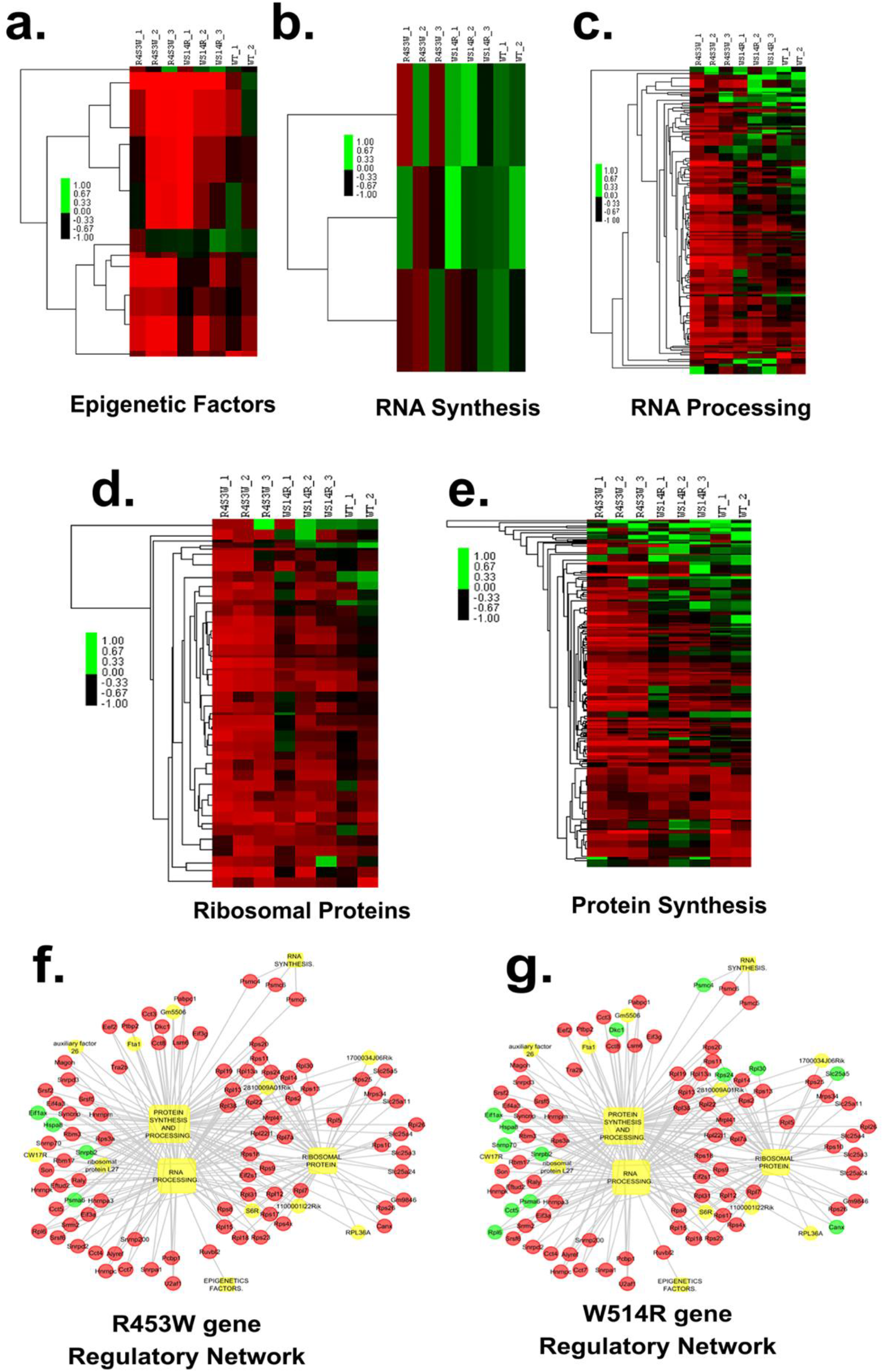
Assessment of individual dysregulated GO categories related to gene expressionregulatory process in mutants. (a), (b), (c), (d), (e) represent heatmaps of epigenetic factors, RNA synthesis, RNA processing, ribosomal proteins, protein synthesis & processing. (f), (g) protein association clusters with proteins name in mutant samples, R453W LA and W514R LA respectively. Red color indicates upregulation and green color indicates down regulation.

Adam et al. (2001) suggested that H2afz protein regulates the recruitment of RNA Pol II enzyme to gene promoters and acts as an important cofactor for the expression of certain genes. Interestingly, we observed a high expression level of H2afz protein in the mutants, which correlates positively with histone activating marks and inversely with the level of H3K27me3 (Jeronima C et al., 2013). Additionally, we found elevated expression of Hist1h1d protein in the mutants (Figure 8), which has been associated with low H3K27me3 modification according to Tiberi et al. (2015). Their study demonstrated enrichment of H3K27me3 levels at the Hist1 locus using ChIP sequencing analysis, and high expression of HIST1 genes in the “low H3K27me3 level” group via qPCR analysis.

**Figure 8.**
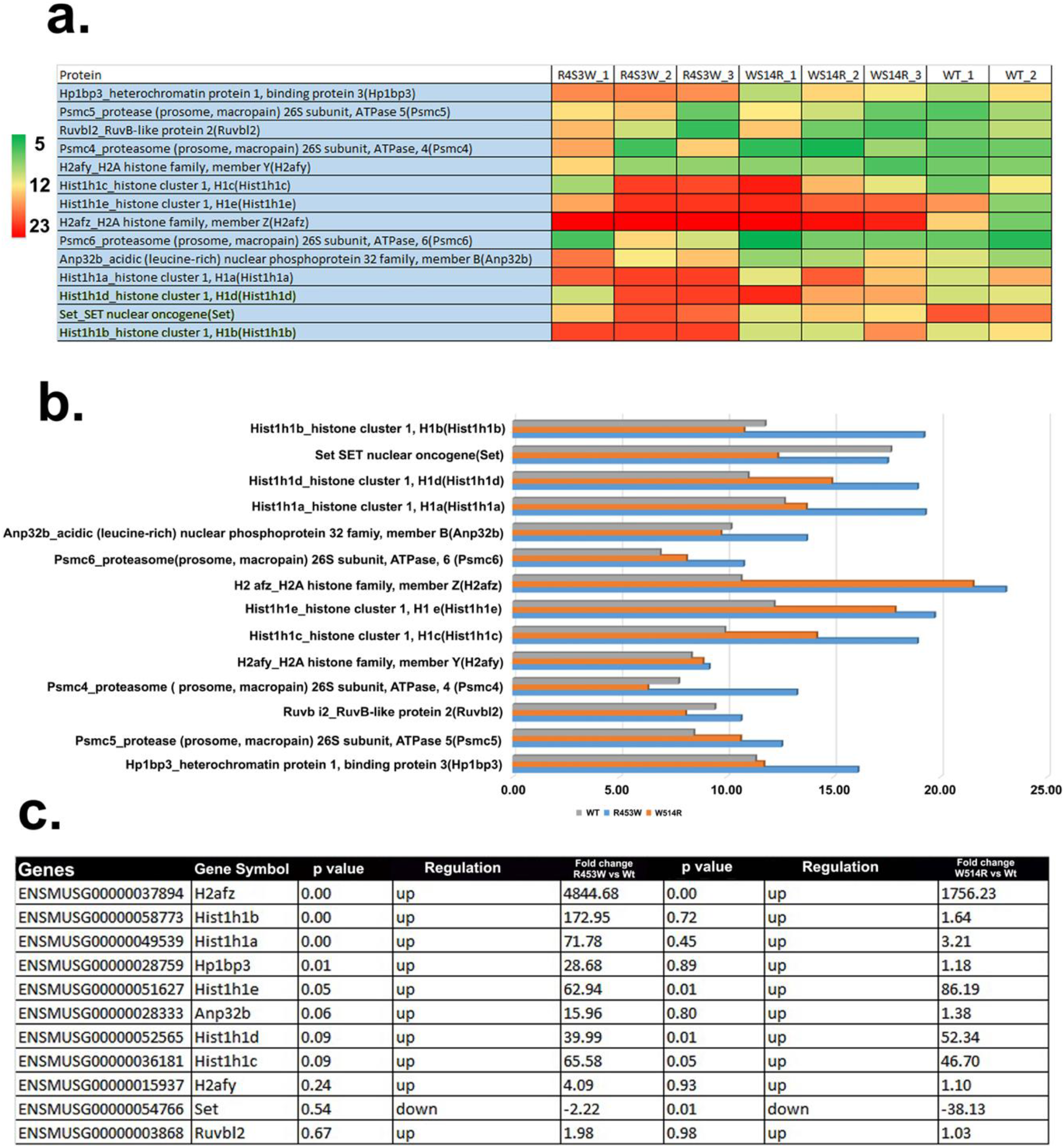
(a) Heatmap of epigenetic regulators within the replicates of 3 samples. (b) Absolute protein level comparison histograms of 3 samples. (c) List of differentially expressed epigenetic regulators in R453W mutant compared to WT LA, W514R mutant compared to WT LA.

Furthermore, we observed high levels of HP1BP3 in R453W mutants (Figure 8), indicating a specific chromatin state, as Garfinkel et al. (2015) reported HP1BP3 co-localization with HP1 dots. Importantly, the altered balance between repressive and activating methylation was confirmed by the decreased expression of SET domain proteins responsible for epigenetic marks in mutant cells.

In our gene regulatory network analysis, we identified upregulation of proteins involved in RNA synthesis, RNA processing, ribosomal proteins, and protein synthesis in the mutants, highlighting significant dysregulation in gene expression regulation. Recent studies have linked some hnRNPs and other splicing factors to lamin A (Roux et al., 2012). Interestingly, we found hyperexpression of members of the heterogeneous nuclear ribonucleoprotein (hnRNP) family in mutant cells, which are involved in pre-mRNA transcription, processing, and availability for other RNA processing factors (Ko et al., 2000; Martinez-Contreras et al., 2007).

### Identification of changes in proteomic profiles that are associated with lamin protein

Dittmer TA et al. (2014) tested 104 interactors through microscopy and yeast two-hybrid screen assays, finding that 53% of interactors are distributed in the nuclear envelope (NE) and 47% throughout the nucleus. Thus, understanding the levels of mutation-specific lamin A interactors could provide insights into disease-specific mechanisms.Using STRING DB, we identified 65 lamin A interactors, with a significant proportion (49/65) showing no change in expression. Specifically, in the R453W LA sample, 16 proteins were upregulated, while in the W514R LA sample, 7 proteins were upregulated and 9 were downregulated (Table 2). These proteins include emerin, lamin B1, lamin B2, and Sun2, which are critical interactors of lamin A involved in maintaining the structural integrity of chromatin and the gene expression layer (May CK et al., 2018; Demmerle et al., 2012; Pascual-reguant L et al., 2018).

**Table 2:**
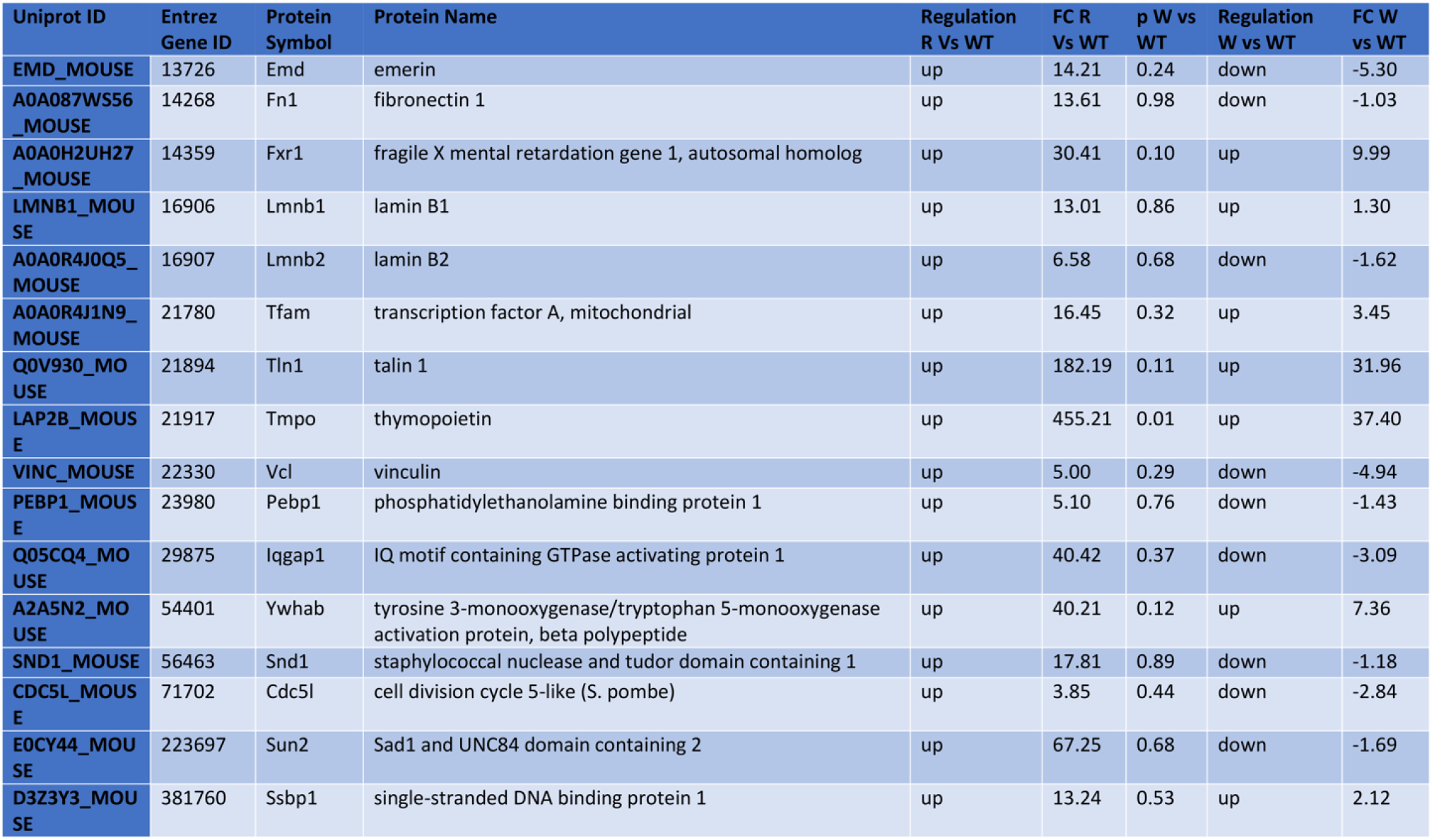
List of differentially expressed protein interactors in R453W mutant compared to WT LA, W514R mutant compared to WT LA.

Understanding the differential expression patterns of these interactors in mutants provides valuable insights into how lamin A mutations may affect nuclear organization and function, contributing to disease pathology.

## Discussion

The nuclear membrane plays a critical role in chromosome organization through various protein-protein interactions, maintaining both peripheral and nucleoplasmic heterochromatin pools (Zuleger et al., 2011, 2013). Mattout et al. (2011) previously highlighted the involvement of heterochromatin in disease mechanisms associated with lamin A mutations. Lamins are believed to interact with chromatin through histones and DNA (Kind et al., 2010). Histone modifications serve as pivotal signatures in gene expression and regulation (Gibney et al., 2010). High levels of H3K9me3 and HP1, along with their dynamic behavior, characterize constitutive heterochromatin (Trojer et al. 2007). However, in mutants examined in this study, there is a notable reduction in H3K9me3 and HP1 marks, suggesting a less condensed chromatin state that may enhance transcriptional activity.Moreover, immunofluorescence and immunoblotting studies indicated a significant decrease in H3K27me3 levels and an increase in H3K36me3 levels. The roles of H3K27me3 and H3K36me3 are complex; while H3K27me3 typically represses transcription, its enrichment around transcription start sites (TSS) suggests a dual role (Young et al., 2011; Shah et al., 2013). H3K36me3 is associated with both constitutive and facultative heterochromatin (Chantalat et al., 2011). Additionally, our proteomics analysis revealed increased expression of Hist1h1d, which correlates negatively with H3K27me3 levels according to Tiberi et al. (2015).

In cells with Hutchinson-Gilford progeria syndrome (HGPS), Scaffidi and Misteli (2006) and Shumaker et al. (2006) observed decreased H3K9me3 levels and increased H4K20me3 levels. The modulation of transcription initiation sites can be influenced by interactions involving epigenetic DNA and histone modifications (Quina et al. 2006). Consistent with this, our immunoblot results indicated increased recruitment of RNA Pol II, potentially leading to enhanced transcription initiation. Surprisingly, our proteomics data showed higher expression levels of regulatory proteins associated with the RNA polymerase II transcriptional preinitiation complex.So, mutations in Lamin A appears to modulate the epigenetic landscape, which could contribute to or underlie the molecular mechanisms of disease pathogenesis.

According to Bank et al. (2011) and Bertrand et al. (2012), the LMNA delK32 mutation is associated with severe early-onset congenital muscular dystrophy, resulting in defective lamin assembly and complete loss of lamin from the nuclear periphery. In Hutchinson-Gilford progeria syndrome (HGPS) cells, expression of truncated Lamin A/progerin leads to loss of H3K9me3, H3K27me3, and HP1, as well as reduction in the peripheral heterochromatin pool (Misteli et al., 2006; Giordano et al., 2014; Lattanzi et al., 2005). Gasser et al. (2011) discovered that retention of muscle-specific promoters at the perinuclear region alters the expression of muscle-specific genes.To relate these findings to the gene expression hypothesis, we can speculate that mutations causing muscular dystrophy lead to lamin A being sequestered into the nuclear interior instead of the periphery, profoundly disturbing the complex interaction between lamin A, lamin-binding proteins, and chromatin at the nuclear periphery. This likely results in broader changes in chromatin organization. Additionally, the localization of nuclear membrane proteins at the inner nuclear membrane depends on their binding to lamins (Ostlund C et al., 2001).

In our study, we observed precipitation of epigenetic marks in nuclear foci associated with these mutations. Previous reports indicate that the nuclear envelope protein Emerin directly interacts with chromatin-modifying enzymes (Holaska et al., 2007), potentially contributing to the observed accumulation of epigenetic marks. However, further research is needed to elucidate how the regulation of these interactions may underlie the reorganization of chromatin states in lamin A-associated muscular dystrophies. The distinct changes in attachment sites and expression patterns observed in our study likely stem from the structural alterations in the lamin A protein due to two different point mutations in the Ig fold domain. Notably, previous research from Bera et al. 2014, Dutta et al. 2018 indicated dissimilar electrostatic surface potentials among these mutants and lower binding affinity towards key nuclear envelope proteins. Specifically, the W514R mutation exhibits an electropositive surface potential in the Ig fold domain, whereas the R453W mutation displays an electronegative surface potential.

In this report, we have gained novel insights into the structure of both mutants. Our rheology experiments revealed that polymer concentration plays a crucial role in influencing the binding strength of these proteins. The W514R mutation exhibited jamming behavior, affecting its load-bearing capacity differently compared to the unjamming behavior observed with the R453W mutation. This observation supports the idea that the lamin intermediate filament acts as a “brick and mortar scaffolding” in re-establishing chromatin interactions with the nuclear envelope. Furthermore, our SEM and 3D SIM images under in vitro conditions indicated differences in network organization between the wild type and the mutants. Complementing these observations, CD spectroscopy experiments confirmed changes in the secondary structure of the overall protein in the mutants. Specifically, W514R lamin A exhibited a structure that aligns more closely with canonical coiled coils. The positive charge in the tail domain of lamin A facilitates crosslinking within the coiled-coil segments, contributing to a spring-like behavior across the entire rod domain.

Therefore, we can extrapolate that the increased positive charge generated by the W514R mutation inhibits flexibility in lamin A and may lead to increased bundling but relatively lower cross-link density in the network, as observed in SEM images. Conversely, the R453W mutation destabilizes the Ig fold domain by disrupting a salt bridge, resulting in a looser network formation by this mutant protein.

Moreover, we observed upregulation of proteins involved in RNA synthesis, RNA processing, ribosomal proteins, and protein synthesis and processing, which may contribute to aberrant splicing, translation, and processing in lamin A-associated muscular dystrophy. This finding supports the notion that the steps involved in gene transcription leading to upstream and downstream translation events are interconnected. It also aligns well with findings by Viera et al. (2014), who reported upregulation of protein translation and RNA processing/splicing in limb–girdle muscular dystrophy. Kong et al. (2010) identified altered splicing of sarcomeric genes such as troponin T2, troponin 13, myosin heavy chain 7, and filamin C gamma, which are crucial biomarkers in dilated cardiomyopathy (DCM) and hypertrophied myocardium, respectively.

In conclusion, our findings suggest a potential link between various types of myopathies characterized by defective RNA and protein processing mechanisms.

## Methods

### Site directed mutagenesis

Site directed mutagenesis was performed using pEGFP –human lamin A,pET-human lamin A constructs (gift from Dr. Robert D.Goldman, Northwestern University, Chicago) following the protocol as described in (Bhattacharjee et al. 2013) to obtain the mutations R453W, W514R. Primer sets each containing the mutations are :

R453W-sense 5’-GGAGGGCAAGTTTGTCTGGCTGCGCAACAA-3’, R453W-antisense 5’-TTGTTGCGCAGCCAGACAAACTTGCCCTCC-3’, W514R-sense 5’-CTAACCGACCTGGTGAGGAAGGCACAGAAC - 3’, W514R-antisense 5’-GTTCTGTGCCTTCCTCACCAGGTCGGTAG - 3’.

### Expression and purification of protein

Full length human lamin A protein/pre-lamin A (664 amino acids) used for this study was expressed from pET-LA, transformed into BL21(DE3)pLysS competent cells and cultured in TB broth (Himedia, Mumbai, India) in the presence of penicillin and chloramphenicol (USB corporation, Cleveland, OH, USA). Protein expression was induced with 2 mM IPTG (Himedia, Mumbai, India) for 2 hours. Cell lysate was prepared as described by Moir et.al. 1991 and separated on a Mono STM5/50 GL Column (GE Healthcare, Uppsala, Sweden) fractions were eluted in 6 M urea, 25 mM TrisHCl pH 8.6, 250 mM NaCl and 1 mM DTT (Urea buffer). Using Slide –A-Lyzer –Minidialysis units with a 10,000 Dalton MWCO(Thermo Scientific, Rockford, IL, USA) proteins were renatured by dialyzing out urea in a step wise manner from 6 M in steps of two at room temperature. 25 mM Tris-HCl buffer of pH 8.6 with 250 mM NaCl and 1 mM DTT, has been used for all experiments with the wild type and mutant proteins. Protein concentrations were determined by standard Bradford reagent (Bio-Rad, Hercules, CA, USA) in a Perkin Elmer Luminescence Spectrometer. Deionized water of highest purity (Resistivity18.2 MV.cm 25°C) obtained from Synergy Millipore water purification system was used for preparing the buffers.

### Scanning Electron Microscopy (SEM)

Renatured protein samples that are in 25 mM Tris-HCl buffer of pH 8.6 with 250 mM NaCl and 1 mM DTT were spotted on circular coverslips (F-13mm), dried in vacuum and coated with gold in IB2 Iron Coater. Samples were imaged in Hitachi S530 Scanning Electron Microscope (Japan) by 3000x magnifications at 25 kV.

### Rheological measurements

The cone and plate rheological measurements were carried out in a stress controlled rheometer (MCR 502, Anton Paar, Graz, Austria) operated through a software (Rheoplus/ 32,service V.3.61). The lower plate is fixed and the shear deformations were applied by rotating the upper cone with a 25 mm diameter, 2°cone -and-plate fixture in a controlled manner. The amplitude (γ) was kept in 1% and angular frequency (ω) was kept in 5 rad/sec‥6 mg/ml of Wild type and mutant proteins in buffer were loaded between the cone-plate and the temperature was maintained at 20°C by Peltier control system. The protein samples were placed in a humidified chamber (with buffer solution) during the measurements, to prevent evaporation of the solvent. The measurements were adequately repeated for 100 measurement points with a duration of 10sec.

### Circular Dichroism (CD) Spectroscopy

Far-UV CD spectra of 4 µM full length human lamin A protein were recorded at 25°C in a Jasco J-720 Spectropolarimeter with a quartz cuvette having a path length of 1 mm. CD spectroscopy were recorded in the far UV range of 200-250 nm at 25°C,with data collected in a continuous mode with a 1nm bandwidth and scan speed of 10 nm/min. Properly refolded proteins were examined after extensive dialysis in 4 M, 2 M urea buffer and finally in the working buffer separately. The spectrum of working buffer at 25°C (average of 10 scans) were subtracted from each spectrum.

### Stable cell line preparation

Mouse myoblast cell line C2C12 was first transfected with EGFP-Wt LA, EGFP-R453W LA, EGFP-W514R LA using Lipofectamine 2000. C2C12 stable cell line were prepared for EGFP-WT LA, EGFP R453W, EGFP W514R using 850 μg of G418 and used in entire cell biology experiments.

### Immunofluorescence analysis

Stably transfected mouse myoblast C2C12 cells grown on 18*18 mm cover glasses were fixed using 4% PFA for 10 min, followed by PBS wash. Cells were permeabilized by 0.2% Triton X-100 solution in PBS stained with antibody anti-Histone H3(tri methyl K27) [Active Motif ; 39155], anti-Histone H3 (tri methyl K9) [Active Motif; 39161], anti-histone (tri methyl K36) [Abcam; ab9050], anti-HP1 alpha [Abcam ;ab234085] of 1:1500 dilution for overnightat 4 C temperature. Then cells were incubated with secondary antibody, Alexa 594 conjugate Invitrogen (A11005) for 2 hour at RT. Finally, coverslips were mounted on Vectashield (Vector laboratories) for imaging.

### Image Acquisition

The slides were visualized in NIKON Inverted Research Eclipse TiE Laser Scanning Confocal/NSIM microscope. Confocal images were captured by Galvano mode NIKON A1 RMP detector and Plan Apochromat VC 100x oil DIC N2 / 1.40 135 NA /1.515 RI objective with an additional 4x digital zoom. A multi-line Argon -Krypton laser (□ ex-457/488/561 nm) at 3% was used for green channel and a solid-state laser (□ ex-561 nm) at 5% was used for red channel. For Z stacks a step size of 0.25 □m was maintained. 25 cells per sample were checked for the study. Every sample was tested for three independent experiments. Images were processed using Ni Elements Analysis Ver 4.13.

3D NSIM images were captured in N-SIM mode using the SR Apochromat TIRF 100×/1.49 NA objective and an iXon3 DU-897E EM CCD camera (Andor Technology Ltd.). The NSIM-488 solid laser line were maintained at 5% of its original power (20 mW). Nuclei were visualized with an additional 4× digital zoom at intervals of 0.15 μm along the z-axis.

For mutant cells, NSIM images were reconstructed with the following parameters: structured illumination contrast set to auto, apodization filter at 0.100, and 3D SIM filter width at 0.05. Images for wild-type LA cells were reconstructed following the protocol described in Shimi et al.2015. All images were acquired using NIS-Elements acquisition software Ver.4.20.00.

### Mathematical equations

To compute elastic energy through mathematical modelling, we can express the co-ordinates i.e. mesh network connecting points as spherical polar co-ordinates (*r, θ, ϕ*) (where r=radial component, *θ*=polar angle, *ϕ*=azimuthal angle) and there is no deformation in the radial part. We can express the deformation in the spherical polar co-ordinates(from r to r + dr, *θ* to *θ* + *dθ, ϕ* to 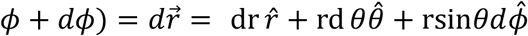. We choose spherical co-ordinates for simplicity in calculation for the shell. We know that Elastic energy (*E*_Total_) of a thin elastic shell can be written as a sum of stretching energy *E*_*S*_ and bending energy *E*_*b*_.i.e *E*_Total_ = *E*_*S*_ + *E*_*b*_ Here we considered that stretching energy is very very greater than bending energy (*E*_*S*_ ≫ *E*_*b*_). Because here we considered a small part in the shell such that the bending energy is very low in that region. So, we can write that the elastic energy = *E*_Total_ = stretching energy = *E*_*S*_. For general case -

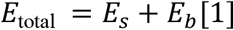

where 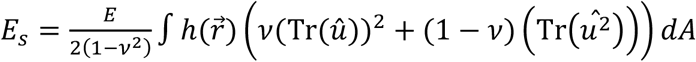and 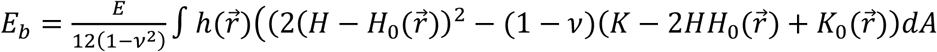

Where H is the mean curvature, and K is the Gaussian curvature; 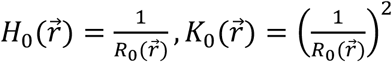 and Gaussian curvatures given in terms of the position-dependent radius *R*_0_ of the reference spherical configuration 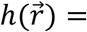 position-dependent thickness of the shell, dA=area element on the surface = *r*^2^sin *θdθdϕ*, E=3D Young’s Modulus,*v*=Poisson’s ratio and 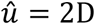 strain tensor with components. We choose *θ* and *ϕ* are the 2 co-ordinates because the deformation happens in *θ* and *ϕ* co-ordinates, while r co-ordinate is fixed.

2 D strain tensor 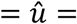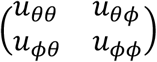 and 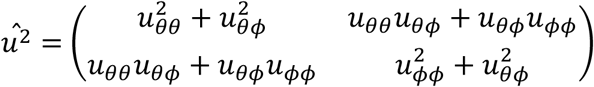

We are interested about the diagonal components because we shall calculate the trace and *u*_*θϕ*_ is symmetric so,*u*_*θϕ*_=*u*_*ϕθ*_.

Where 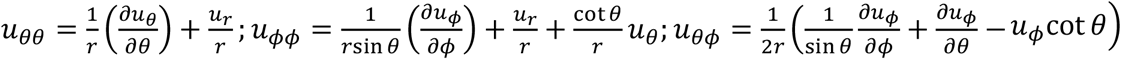

Let’s say 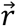 and 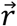 are the positional vectors of mutant nucleus and wild type cell nucleus. We can write the Displacement vector as

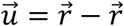 for small changes in displacement vector 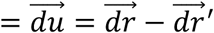 if 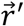 is constant then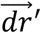 is 0.Then 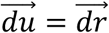

In polar form we can write 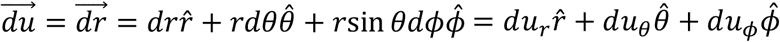 Where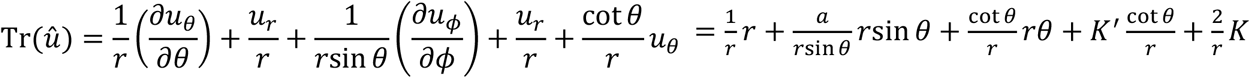 from equation (i).

Here 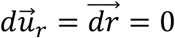

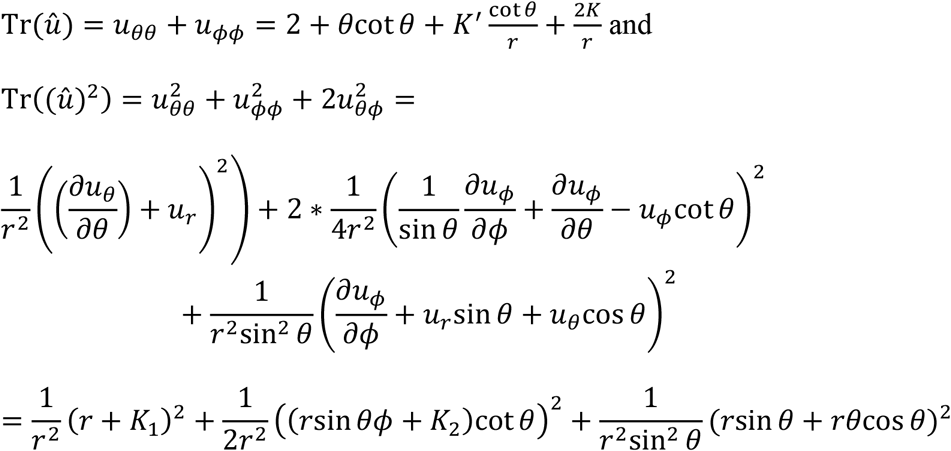

*K, K*^’^, *K*_1_, *K*_1_, *K*_2_ and *K*_3_ are arbitrary integration constants.

So, 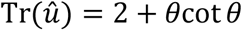 and

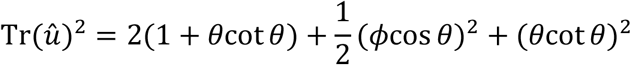

we choose all the arbitrary constants are 0 because the value of constants are really small.

now elastic energy due to nuclear lamin A = Elastic energy due to nucleus (i.e. nuclear envelope proteins) + elastic energy due to nodes (nodes are formed due to nuclear lamin A)+ elastic energy due to mesh of chromatins

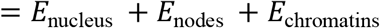

But 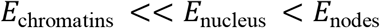 where 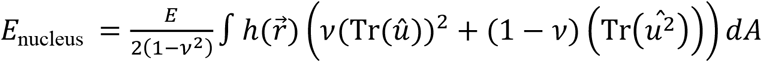 and

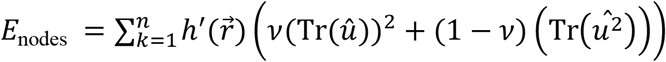

here n is the no. of nodes. Here the no. of nodes depends upon the azimuthal angles and we summed it over *θ* and *ϕ*.Here we calculated upto *θ*=π/2

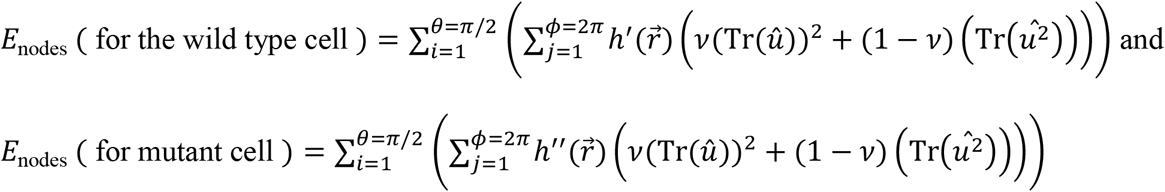

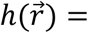Constant for the nuclear envelope ; 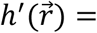 Constant for wild type cells nuclear lamin A nodes ; 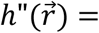 Constant for mutant cells nuclear lamin A nodes.

where Φ=*m* ∗ *ϕ* and Θ=*n* ∗ *θ*.

m and n are the no of slices and no. of points on the slice. The numbers are discrete integers.

### Fluorescence recovery after photo bleaching (FRAP)

C2C12 cells were grown on Nunc Glass Bottom Dish, 27 mm (Cat no.150682) in High Glucose Dulbecco’s modified Eagle medium (DMEM) (Gibco), complemented with 10% fetal bovine serum (Gibco) and 1% penicillin-streptomycin (Gibco), in a humidified incubator at 37°C in 5% CO2. Cells were synchronized at G2/M phase with 200 ng/ml nocodazole (Sigma) for 12 hrs‥ Cells were washed three times with PBS and replenished with pre-warmed fresh complete media after release form the block. After 3hrs from release of the nocodazole block cells were double transfected with pEGFP-human LA (EGFP-wt LA and mutants) and RFP-HP1α using Lipofectamine 2000 (Invitrogen) in accordance with the manufacturer’s protocol at early passages having 60-70% confluency and were kept in culture for 24 hrs.

FRAP experiments were performed on a Nikon Inverted Research Eclipse TiE Laser Scanning Confocal/NSIM microscope. A 2um * 2um region at the nuclear periphery in the mid-focal plane was bleached at 100 % power of the argon laser (405 nm) running at 40.6 mw. The pinhole size for the confocal was set at 1 Airy unit. Bleaching was done for 5s followed by 50s for recovery. 30 cells per sample was selected for FRAP analysis from at least 2 independent experiments. Immediately, following the bleach, images in the Z plane were captured. The best Z-plane image for the bleached ROI was selected from each time point 3D stack to avoid cell movement, nuclear rotation and focus drift to register the bleach ROI between time points. Cells were maintained at 37. C,5% CO_2_ using a steam incubator. Data was normalized using following equation:

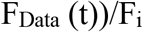

F_Data_(t) = Background corrected FRAP data at specific time point.

F_i_ = Prebleach steady state fluorescence.

We have used following equations to calculate the halftime (t_1/2_) of recovery and mobility fraction (M_f_) :

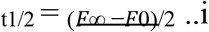

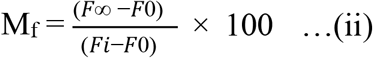

### Immunoblotting

Stably transfected cells were harvested after rinsing off the media with 1x PBS. Cells were lysed in RIPA buffer with 7M Urea (50mM Tris-HCL pH 8.0, 150 mM NaCl, 0.75% SDS (w/v), 0.5% Sodium Deoxycholate (w/v), 1% Triton X-100 (v/v), on ice for 30 min followed by sonication with UP200S Ultra Schallprozessor at 0.5 cycles, amplitude set at 60-70% to lyse the cells until the solution become transparent. Cell lysate was mixed with chilled .acetone at 1: 3 ratio and kept for overnight at – 20^·^ C. On the next day cell pellet was achieved by centrifugation at 13000 rpm for 10 min and resuspended in RIPA buffer with 7M Urea. After protein estimation by Bradford reagent using Nanodrop 2000 spectrophotometer (Thermo scientific) and equal amounts of protein (20 ug)were loaded and resolved on 15% SDS PAGE.Proteins were transferred to nitrocellulose membrane, and probed as with primary (overnight at 4 °C) and secondary antibodies (2 h at RT), Following antibodies were used :

| Antibody | Source | Catalog number | Dilution |
| --- | --- | --- | --- |
| Histone H3 (tri methyl K9 ) | Active Motif | 39161 | 1:1000 |
| Histone H3 (tri methyl K27 ) | Active Motif | 39155 | 1:1000 |
| Histone H3 (tri methyl K36 ) | Abcam | ab9050 | 1: 1000 |
| HP1 alpha | Abcam | ab234085 | 1:800 |
| RNA polymerase II<br>CTD (4H8) | Abcam | ab5408 | 1: 1000 |
| $\beta$ Actin | Sigma–Aldrich | A5316 | 1:800 |

Super Signal West Pico Chemiluminescent Substrate (Thermo Scientific) was used to develop chemiluminescence and imaged using Kodak Medical X-ray Films. Blots were repeated in triplicates if not otherwise mentioned. Quantifications from the blots were performed using Image J software. Data presented as bar diagrams by normalizing the intensity of histone modification marks, RNA Pol II S5p marks with loading control β actin.

### LC-MS/MS

20 ug of protein per sample from three biological replicates per treatment group and two from control was taken for analysis. Sample were alkylated in dark for 20 mins at RT with 30 mM Iodoacetamide following reduction with 10m mM DTT at 56 ° C for 30 mins. pH of the solution was adjusted to 8 by adding 25 mM of ammonium bicarbonate. Then trypsin (0.2 μg/μL) was added to the solution so that the concentration of trypsin: proteins was 1: 50. After trypsin addition, the samples were incubated overnight at 37° C. 5 % formic acid was added to stop the reaction. Chromatographic separation was achieved on Pep map TM 100(75 μmx * 2 cm); Nanoviper C 18, (3 μm; 100 A°) as precolumn and EASY SPRAY PEPMAP RSLC C18 2 μM (50 cm * 75 μm;100 A°) as analytical column. .4 μg sample was injected onto column. The mobile phase A was 0.1% Formic acid in MS grade water and mobile phase B was 80% Acetonitrile + 0.1% formic acid in MS grade water. Step gradient Mobile phase B was used from 10% to 95% for 3 hr with 300 μL/min flow rate. A survey full scan MS (from m/z 375-1700) was acquired in the Orbitrap Fusion detector with a resolution of 120000. MS/MS was performed with a scan range m/z 100-2000 in the Ion trap detector. Protein identification was performed by searching MS/MS data against the swiss-prot mouse protein database downloaded on Oct 2, 2018 using the in house mascot 2.6.2 (Matrix Science) search engine. The search was set up for full tryptic peptides with a maximum of two missed cleavage sites. Acetylation of protein N-terminus and oxidized methionine were included as variable modifications and carbamidomethylation of cysteine was set as fixed modification. The precursor mass tolerance threshold was set 10 ppm for and maximum fragment mass error was 0.02 Da. The significance threshold of the ion score was calculated based on a false discovery rate of ≤ 1%. Qualitative analysis was performed using progenesis QI proteomics 4.1 (Nonlinear Dynamics).

### Statistical Analysis

Data are presented as means ± SD. The entire significance test for immunofluorescence and immunoblot assays were obtained using student’s t-test. All the experiments were repeated for three times.

Origin (OriginLab, Northampton,MA) was used for all FRAP analysis.

The effect of mutations on the structural stability was predicted With DynaMut2.

## Acknowledgments

The authors thank Dr. Dipayan Sanyal (CSIR-CGCRI), Dr. Srikanta Chakraborty (University of Burdwan) for sharing of equipments and insightful discussion. For LC-MS/MS authors acknowledge Biologics Characterization Facility, C-CAMP. We thank Dr. Prabal Chakraborty, application Scientist, Towa Optics India Pvt Ltd. for valuable inputs regarding image processing. We are grateful to Dr. Alejandro Vaquero (Josep Carreras Leukemia Research Institute, Spain) for RFP HP1α construct. This research was partly funded by SERB-DST and BARD project,DAE.

## Author Contributions

S.D.,M.V. conceived and designed the experiments. SD Performed all the immunofluorescence, immunoblot studies and biophysical experiments. S.D., M.V. and V.K. analyzed the proteomics data. A.B done the mathematical modelling. S.D., M.V. made the draft of the manuscript. S.D edited the manuscript. All authors have read and agreed to the published version of the manuscript.

## Data availability

All relevant data, associated protocols, and materials are within the manuscript and its supplemental material. Corresponding authors are willing to make available additional information upon reasonable request.

