## Supplementary material for "Investigating the differential structural organization and gene expression regulatory networks of lamin A Ig fold domain mutants of muscular dystrophy": NA

**Figure S1**

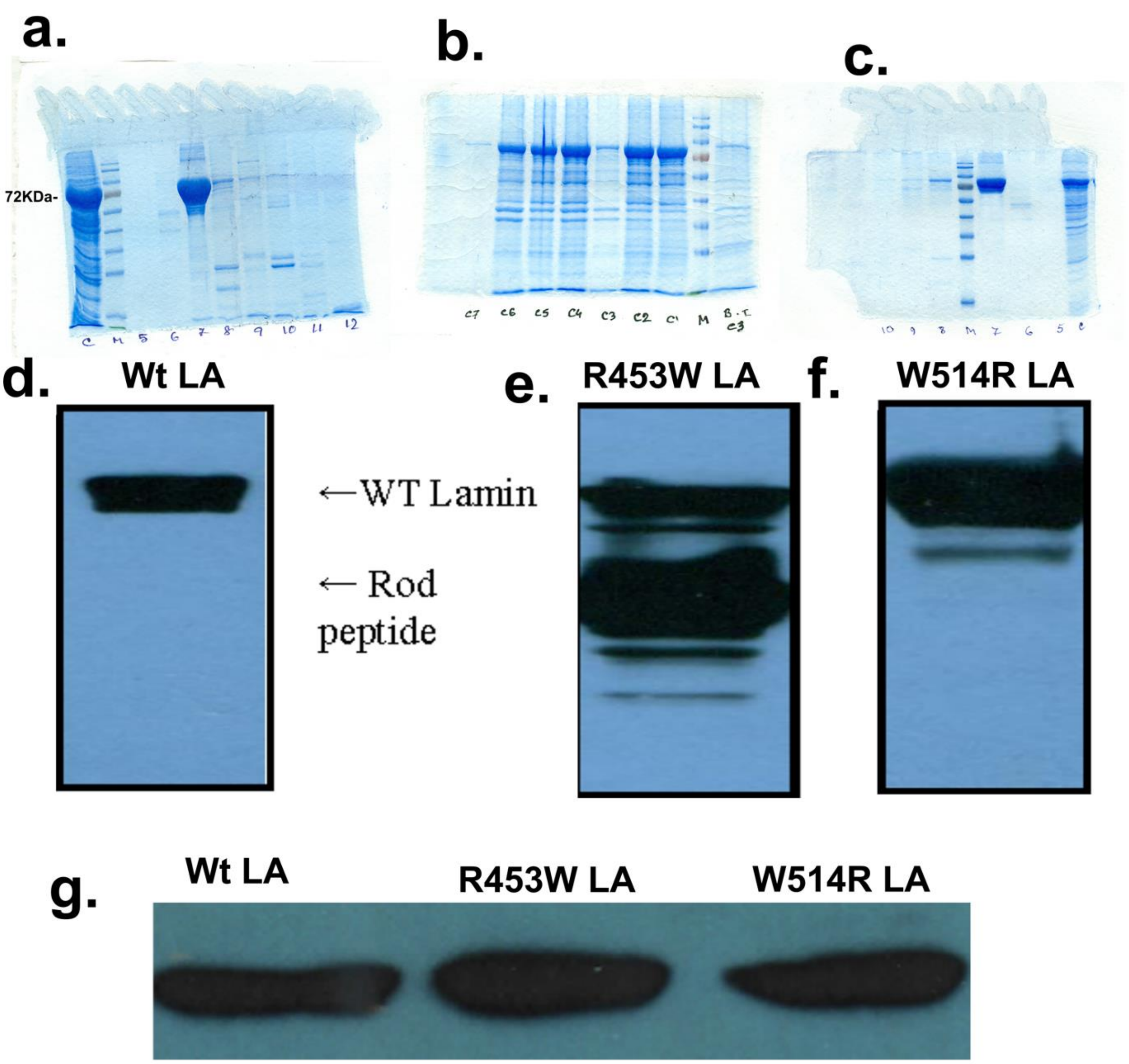


**Figure S1: (a),(b) and (c) present 10% SDS PAGE of protein expression of Wt LA ,R453W LA, W514R LA proteins .At analytical level R453W shows degradation of the protein in rod peptide. Fractions 5th -12th were collected from column purification and loaded from lane 3-10.Lane 1 -crude protein,2-marker, maximum protein for Wt LA found in lane 5,for R453W lane 3-6 (starting from right) contain protein samples and for W514R lane 4 shows maximum protein samples of fraction 7 .**

**(d),(e), (f) blots of Wt LA and mutant LA s - using JOL2 primary antibody after first round of column purification.(g) blot represents purity of protein samples after final round of column purification.**

**Figure S2**

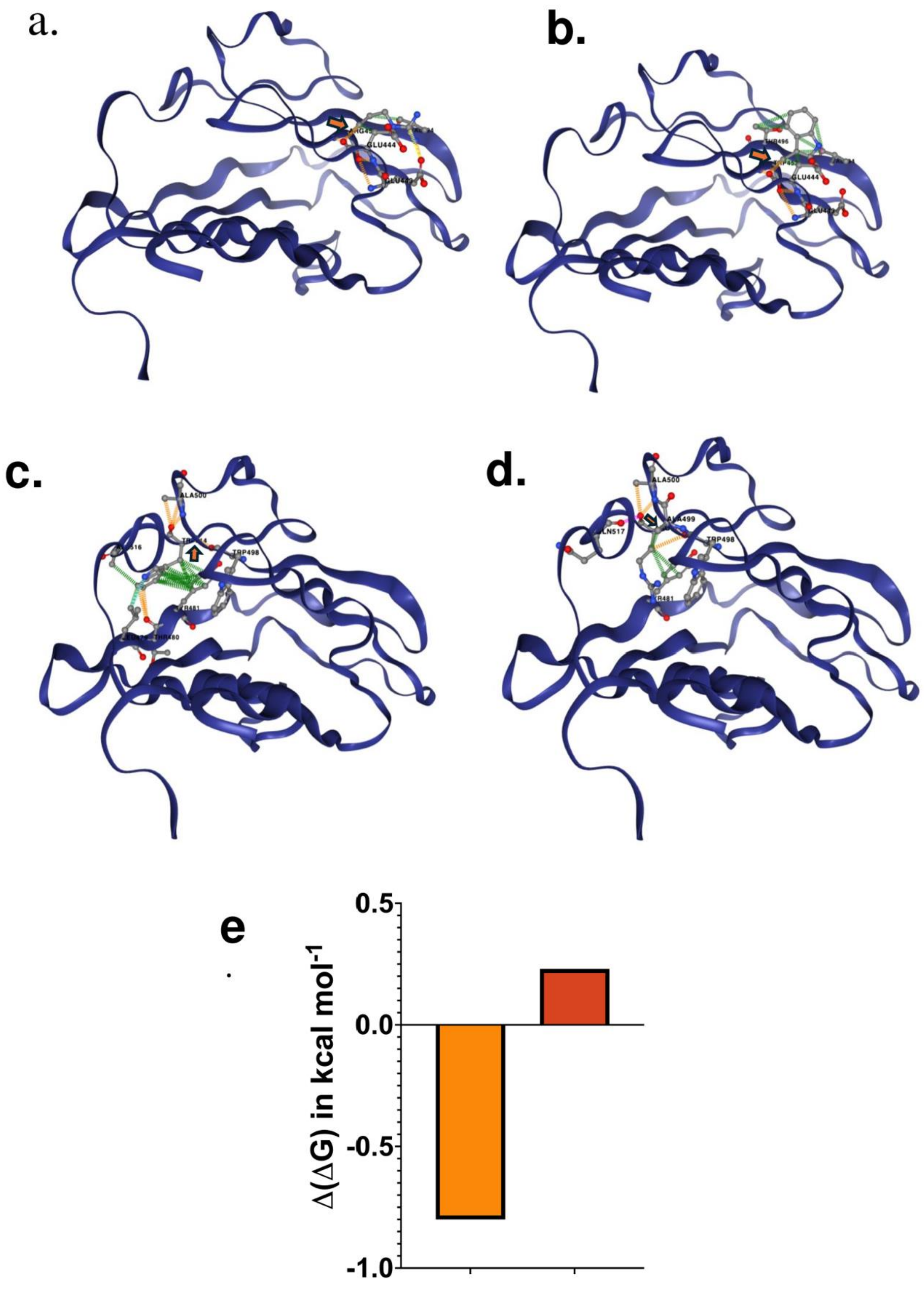


**Figure S2 :** Changes in protein stability with Arg 453-Trp and Trp 514-Arg (a) represents

R453 - Ig Lamin A fragment (PDB ID : 1 IVT) and (b) depict W453- Ig Lamin A fragment. ( c) and (d) are cartoon representations of Trp 514-Arg Ig Lamin A fragment.
(e) DynaMut2 server was used to predict the change in the free energy of unfolding (ΔΔG) induced by the mutations Arg 453-Trp and Trp 514-Arg . Arg 453-Trp has a negative ΔΔG while Trp 514-Arg shows positive ΔΔG .

**Figure S3**

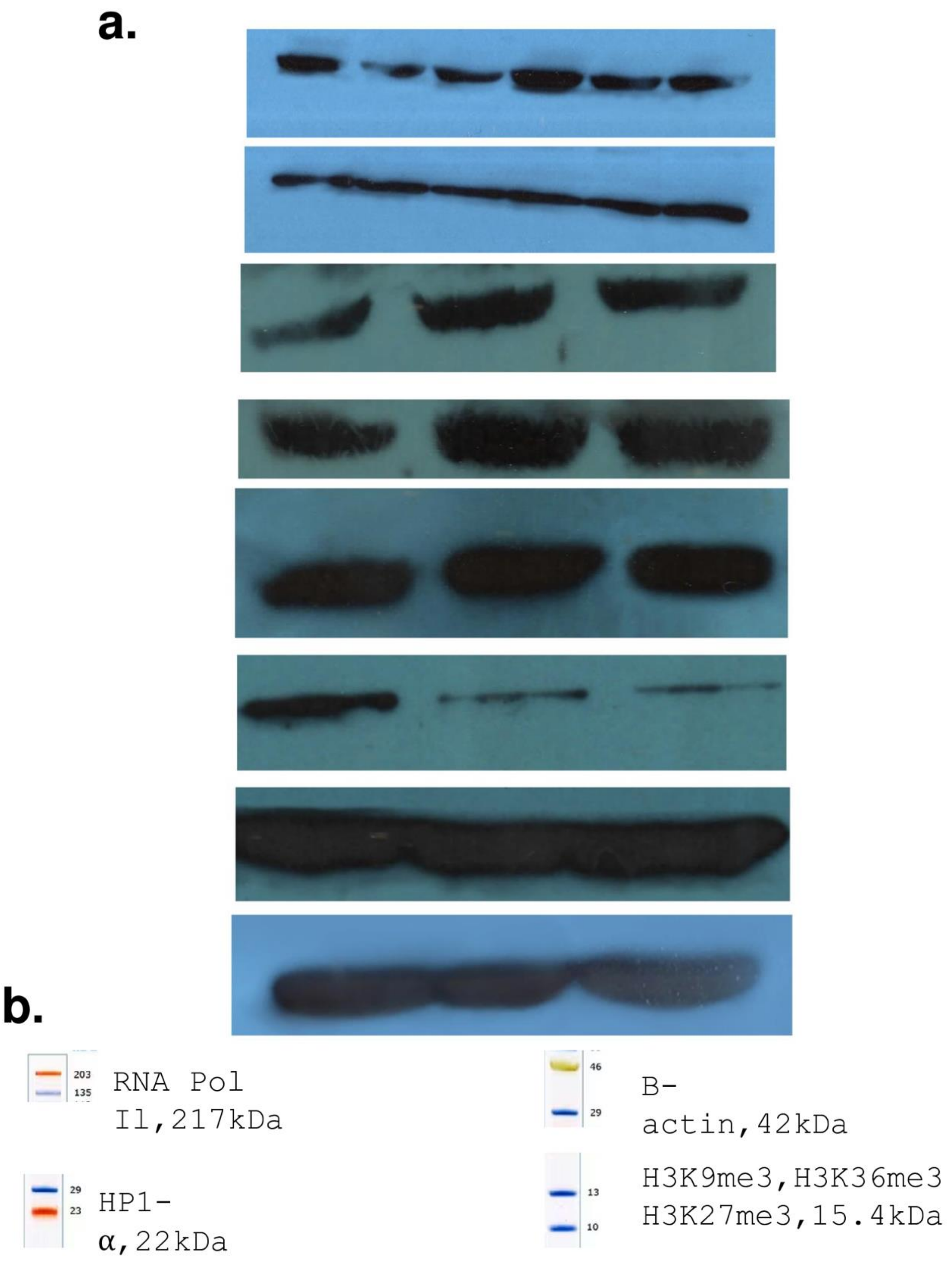

**Figure S3:** (a.) represents raw blot images used in main file.

(b) suggests cutting positions of .45 um membrane according to the molecular weight of targeted proteins.

**Figure S4**

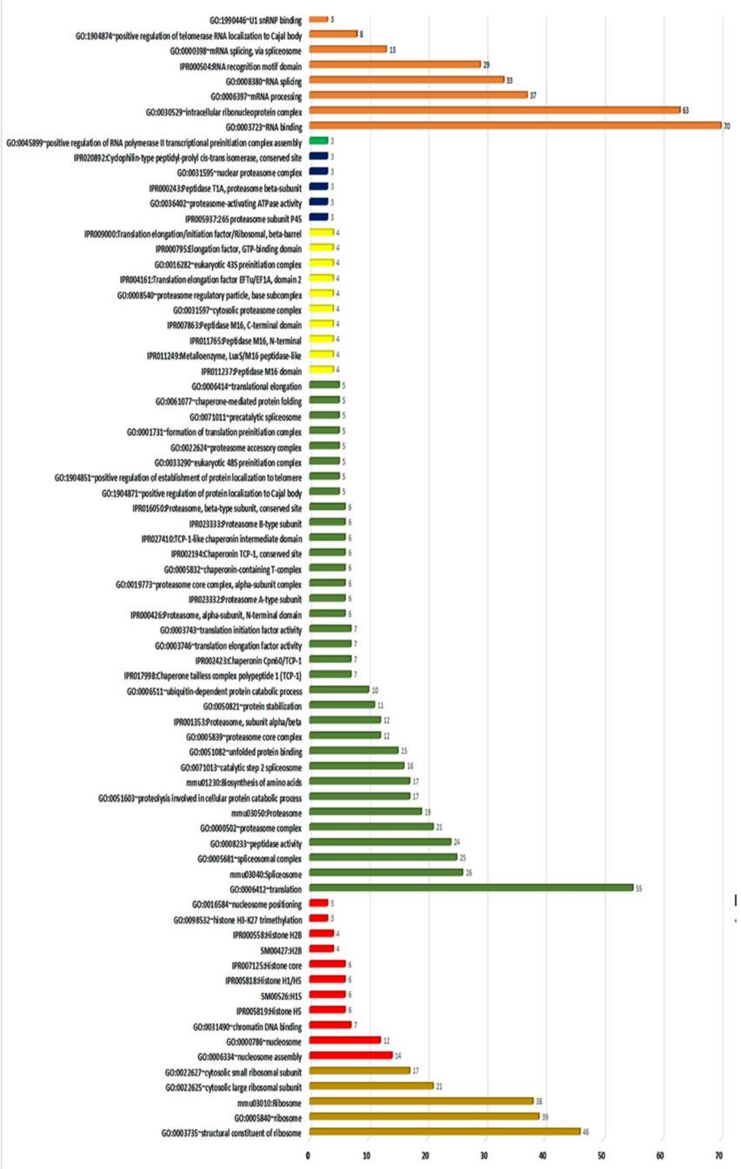

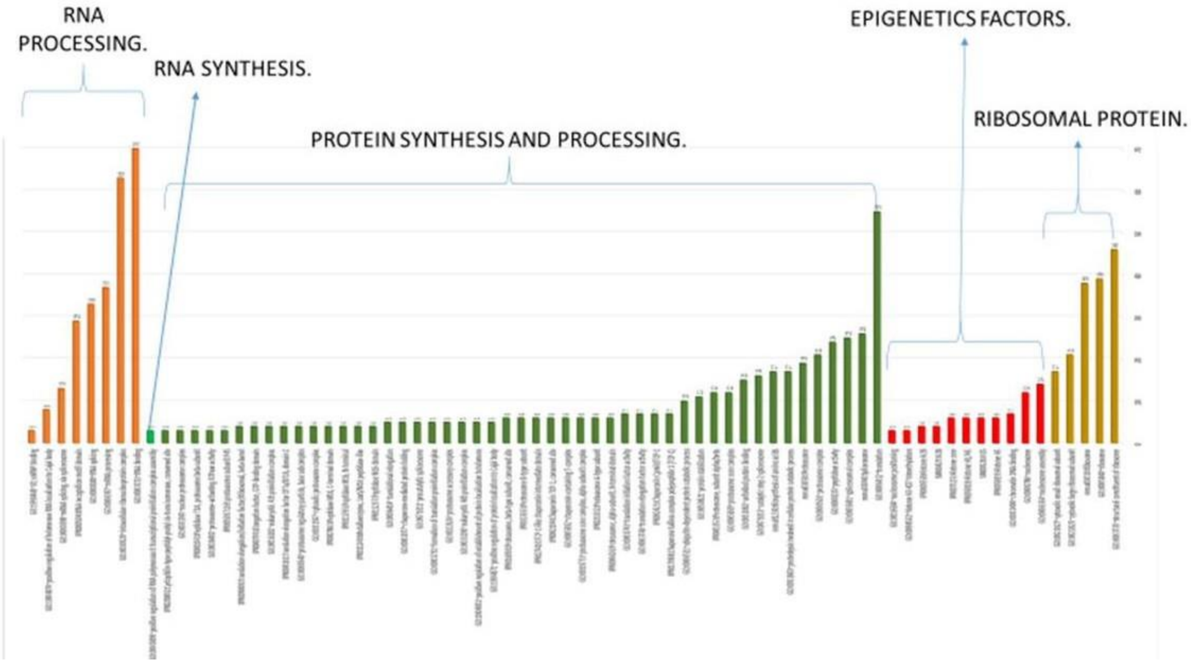


**Figure S4:**

**Histograms of dysregulated GO categories related to gene expression regulatory processin mutants.**
